# Hidden in plain sight: The effects of BCG vaccination in COVID-19 pandemic

**DOI:** 10.1101/2020.06.09.142760

**Authors:** Eman Ali Toraih, Jessica Ashraf Sedhom, Titilope Modupe Dokunmu, Mohammad Hosny Hussein, Emmanuelle ML Ruiz, Kunnimalaiyaan Muthusamy, Mourad Zerfaoui, Emad Kandil

## Abstract

To investigate the relationship between BCG vaccination and SARS-CoV-2 by bioinformatic approach. Two datasets for Sars-CoV-2 infection group and BCG-vaccinated group were downloaded. Differentially Expressed Genes were identified. Gene ontology and pathways were functionally enriched, and networking was constructed in NetworkAnalyst. Lastly, correlation between post-BCG vaccination and COVID-19 transcriptome signatures were established. A total of 161 DEGs (113 upregulated DEGs and 48 downregulated genes) were identified in the Sars-CoV-2 group. In the pathway enrichment analysis, cross-reference of upregulated KEGG pathways in Sars-CoV-2 with downregulated counterparts in the BCG-vaccinated group, resulted in the intersection of 45 common pathways, accounting for 86.5% of SARS-CoV-2 upregulated pathways. Of these intersecting pathways, a vast majority were immune and inflammatory pathways with top significance in IL-17, TNF, NOD-like receptors, and NF-κB signaling pathways. Our data suggests BCG-vaccination may incur a protective role in COVID-19 patients until a targeted vaccine is developed.

**Supplementary Materials:** **(https://drive.google.com/open?id=15Na738L282XNaQAJUh0cZf1WoG9jJfzJ)**

## INTRODUCTION

The coronavirus pandemic now affects over 2 million people with nearly 130,000 deaths reported thus far, resulting in unprecedented ramifications on global health, social infrastructure, and economic trade. Since its emergence in Wuhan, China in 2019, SARS-coronavirus-2 (SARS-CoV-2), the causative agent of the disease, has spread rapidly across the globe to over 185 countries. Curiously, the degree of intensity has varied markedly between countries - even those with similar climates, geography, and/or healthcare infrastructure. The transmission pattern does not appear to follow previous SARS-CoV virus transmission^1^ or climatic zones based on human transmission^2-4^ as temperate and tropical countries have reported the disease^5^. The incidence of COVID-19 pandemic varies widely with the highest number of cases in the United States, Italy, China and lowest in regions of the world historically vulnerable to infectious disease such as sub-Saharan Africa. The disproportionately high morbidity and mortality of COVID-19 infection in some countries has been linked to several factors including, but not limited to differences in efforts to mitigate the disease spread (i.e. social distancing, travel restrictions), circulating SARS-CoV strains, and vaccination policies.

In order to confront COVID-19 disease, it is critical to characterize the virus’ biological and immunological pathways. Recent studies have uncovered the mechanism of viral entry of the SARS-CoV-2 relies on human angiotensin converting enzyme 2 (hACE2) acting as the host cell receptor. hACE2 interacts with the glycoprotein spike (S) of SARS-CoV2 in order to form a complex structure, a receptor-binding mode motif similar to some related viruses, including the one responsible for Severe Acute Respiratory Syndrome (SARS)^6-11^. Another study, which analyzed the transcriptional changes in the immune genes of 3 COVID patients, reported increased inflammatory responses to the virus, revealing increased T-cell activation and cytokine expression, indicating pro-inflammatory pathways may be prognostic markers and/or serve as potential targets in COVID-19 disease^12^. Increased IL-6 expression and tumor necrosis factor (TNF-α) has also been reported, furthering the notion that cytokines are playing a key role in COVID-induced pneumonia^13^. Other studies have hypothesized that protection by the Bacille Calmette-Guérin (BCG) vaccination, which was originally developed to protect against Tuberculosis (TB) disease, may be playing a critical immunological role against CoV-Sars-2^14-19^.

The BCG is a century-old vaccine that is given as an attenuated live strain of *Mycobacterium bovis* used to confer immunity against some strains of TB. Various studies have indicated that BCG vaccination has a role extending far beyond TB treatment alone, eliciting non-specific effects (NSEs) within the innate immune system alongside the adaptive immune response^20^. BCG vaccination has been identified for its protective role specifically against respiratory viral infections, including Influenza A and respiratory syncytial virus (RSV)^21^, and is the standard therapy for certain types of bladder cancer. Although the exact mechanism remains elusive, BCG-protection is thought to be conferred by epigenetic and immunological moderation of the immune response through the release of pro-inflammatory cytokines (TNF-α, IL-6, and IFNγ) and the role of Vitamin D^22-24^. One such proposed mechanism behind the protective effects of BCG-vaccination heavily implicates the role of Vitamin D. During antimicrobial response (such as in *M.tb* infection), toll-like receptors (TLRs) upregulate the expression of VDR on immune cells and pulmonary epithelial cells. VDR then binds to calcitriol, the active form of Vitamin D, and together regulates the transcriptional activity of several antimicrobial peptides such as cathelicidin and beta-defensin^25-28^. Vitamin D alters T-cell activation and IFN-gamma stimulation, increasing the expression of several pro-inflammatory cytokines. Overall, the non-specific effects following BCG vaccination are conferred by epigenetic and transcriptional modulation of the innate immune system as evidenced by its far-reaching role in several viral infections.

Furthermore, it has been hypothesized that countries with nationalized BCG vaccination policies show decreased morbidity and mortality to COVID-19 when compared to those where no such uniform policy exists (such as the United States or Italy); however, these preliminary data is limited given it is yet to be peer-reviewed and fails to account for several confounding factors such as age, testing rates^29,30^, and the accuracy of the BGC World Atlas^31^(http://www.bcgatlas.org/). Nonetheless, given the safety of BCG vaccination and the well characterized role of NSEs, it is theorized that BCG can serve as a temporary and safe solution until a targeted vaccination becomes available. Currently, four clinical trials are already underway in Australia, Netherlands, and the United States involving BCG vs placebo-controlled trials in health care workers involved in Covid-19 patient care (ClinicalTrial.gov; *NCT04347876, NCT04327206, NCT04328441, NCT04348370*). Further, there is an additional observational study in Egypt for tuberculin positivity in COVID-19 patients vs COVID-19 negative, BCG-vaccinated parallel cohort (*NCT04350931*).

Overall, the far-reaching effects of BCG vaccination testify to its dynamic role: eliciting NSEs, curbing inflammation in cancer models, and reducing viremia in several distinct pathogens, including RSV, Influenza A, yellow fever, and herpes simplex virus^21,32^. Our inclination is that BCG vaccination may provide putative immunity against COVID-19 until a targeted vaccine can be well studied and mass produced. Altogether, this study aims to elucidate the relationship between BCG vaccination and SARS-CoV-2 through bioinformatic analysis of biological and immunological pathways underlying both.

## MATERIALS AND METHODS

### Data Source

The microarray data analyzed in this study were obtained from the Gene Expression Omnibus (GEO) (https://www.ncbi.nlm.nih.gov/geo/), accession number GSE147507, published on March 25^th^, 2020. This data set analyzed gene expression profile of a normal human bronchial epithelial (NHBE) cell line, derived from a 79-year-old Caucasian female, after SARS-CoV-2 viral infection. The following samples were analyzed; SARS-CoV-2 infected NHBE cells (GSM4432381, GSM4432382, GSM4432383) compared to mock treated NHBE cells (GSM4432378, GSM4432379, GSM4432380).

To compare the transcriptomic alterations in SARS-CoV-2 viral infection with host immune response following administration of BCG vaccine, the RNA sequencing data (GSE87186) was retrieved from the SRA database (https://www.ncbi.nlm.nih.gov/geo/query/acc.cgi?acc=GSE87186) in Biojupies Analysis Notebook (https://amp.pharm.mssm.edu/biojupies/analyze) and processed by ARCHS4 (all RNA-seq and ChIP-seq sample and signature search) pipeline (https://amp.pharm.mssm.edu/archs4)^33^. In this dataset, the immunogenicity of a recombinant BCG vaccine in healthy BCG-naïve adults and who were negative for prior exposure to *M.tb* was evaluated. We analyzed 40 samples of 8 adult vaccine recipients at 5 different timing post-vaccination; representing day 0, day 14, day 28, day 56, and day 84 (GSM2324141-GSM2324180). Separate analysis was first performed followed by combined analysis.

### Data pre-processing and differential expression analysis

After background correction of raw expression data, mapped multiple probes to the same genes were summarized to gene levels by using the median. Genes were filtered out if variance percentile rank lower than the threshold (set at 15%) or low relative abundance (average expression signal) below 5%. To ensure similar expression distributions of each sample across the entire experiment, normalization by log2 transformation method was performed. The quality of the normalized dataset was checked with the box plot and density plot, and the Benjamini and Hochberg method was selected for the multiple testing correction. Using the Limma R package available on Bioconductor (http://bioconductor.org/packages/release/bioc/html/limma.html), differentially expressed genes (DEGs) were identified in the microarray dataset. Genes with false discovery rate (FDR) <0.05 and log_2_-fold change (FC) ≥1.0 were considered as significantly differentially expressed and subjected to further analysis. For the RNA seq data of BCG vaccine, gene expression signature was generated by comparing gene expression levels of the post-vaccination groups with the control group at day 0. Altered patterns of gene expression were defined through each experimental time (FC>1|P<0.05). To visually identify the similarities and differences between different cell line samples, principal component analysis (PCA) was plotted to project high-dimensional data into lower dimensions using linear transformation.

### Functional Enrichment Analysis

Gene Ontology (GO) database for function annotation of genes was used to analyze the DEGs at the functional level in terms of biological processes, molecular function, and cellular component domains^34^. Pathway functional analysis was performed in the Kyoto Encyclopedia of Genes and Genomes (KEGG) database, Wikipathway, and reactome pathways ^35^. The significantly overrepresented GO and pathways of the up-regulated and down-regulated DEGs were identified by The Search Tool for the Retrieval of Interacting Genes (STRING) database version 11.0 (https://string-db.org/) and validated in Enrichr website (http://amp.pharm.mssm.edu/Enrichr/)^36^ with p value <0.05 as a cut-off criterion.

### Protein-protein interaction network construction

The STRING database was used to construct protein interaction pairs of the screened DEGs based on their function and scores. Setting was adjusted with high confidence (combined score) >0.9 and existence of experimental evidence. The network was visualized in Network Analyst (https://www.networkanalyst.ca/), a visual analytics platform for comprehensive gene expression profiling with the following threshold; degree distribution of 20 and betweenness at 10. The hub nodes in the protein-protein interaction (PPI) network were then identified based on the connectivity degree in the network statistics (number of neighbors).

### Gene regulatory networks

Gene-miRNA Interactome was constructed from experimentally validated miRNA-gene interaction data collected from TarBase v7.0 (http://diana.imis.athena-innovation.gr/) and miRTarBase (http://mirtarbase.mbc.nctu.edu.tw/) databases and plotted in Network Analyst. Potential transcription factors (TFs) among DEGs were first screened in the Panther (http://pantherdb.org/), JASPAR TF binding site profile database (http://jaspar.genereg.net/) and the ENCODE ChIP-seq data (https://www.encodeproject.org/chip-seq/transcription_factor/). In addition, upstream regulatory networks for the DEGs were built, and inferred networks combining transcription factor enrichment analysis and protein-protein interaction network expansion with kinase enrichment analysis was constructed in Expression2Kinase web application (http://amp.pharm.mssm.edu/X2K/).

### Correlations between post-vaccination host immune response and COVID transcriptome signatures

Through KEGG pathway enrichment analysis, down-regulated KEGG signaling pathways following BCG vaccination were intersected with up-regulated pathways in SARS-CoV-2 infection using Venny 2.1.0 (https://bioinfogp.cnb.csic.es/tools/venny/). Deregulated genes in each pathway were explored and the direction of expression were compared between SARS-CoV-2 and BCG-vaccinated experiments in KEGG mapper (https://www.genome.jp/kegg/mapper.html).

## RESULTS

### Transcriptomic changes in SARS-CoV-2 infection

#### Data exploration

Raw data for the reads is provided in **Supplementary Table S1**. After normalization, the quality was checked in density and box plots (**Supplementary Figure 1A, B**). Mock-treated and infected samples showed clear discrimination pattern in PCA, **Supplementary Figure 1C**.

#### Differentially expressed genes screening

Of 23710 genes screened in the experiment, 7107 genes with constant values were removed. A total of 161 annotated genes were identified to be differentially expressed, which included 113 upregulated DEGs and 48 downregulated genes (**Supplementary Figure 1D**). The top variable genes between the SARS-CoV-2 and mock treated samples is shown in **Figure 1**. Of the up-regulated genes with the highest expression level, (a) Small Proline Rich Protein 2F (*SPRR2F*) which is an important structural protein that provides a protective barrier in stratified squamous epithelium and is associated with anti-microbial response; (b) Granulocyte-colony stimulating factor (*CSF3*), a proinflammatory cytokine, has been shown to impair CD8+ T cell functionality and act as a modulator of T-Cell and dendritic cell functions, (c) Intercellular adhesion molecule 2 (*ICAM2*) is expressed on bronchial epithelial, mediates adhesive interactions important for antigen-specific immune response, NK-cell mediated clearance, and lymphocyte recirculation, (d) *S100A7A*, immunogenic-related calcium binding protein, regulated by TLR4, and associated with psoriasis, (e) TNFAIP3-interacting protein 3 (*TNIP3*), which binds to zinc finger protein and inhibits NF-kappa-B activation induced by tumor necrosis factor, Toll-like receptor 4 (*TLR4*), and (f) *PCNA-AS1*, the antisense and regulator for Proliferating Cell Nuclear Antigen gene.

**Fig. 1.**
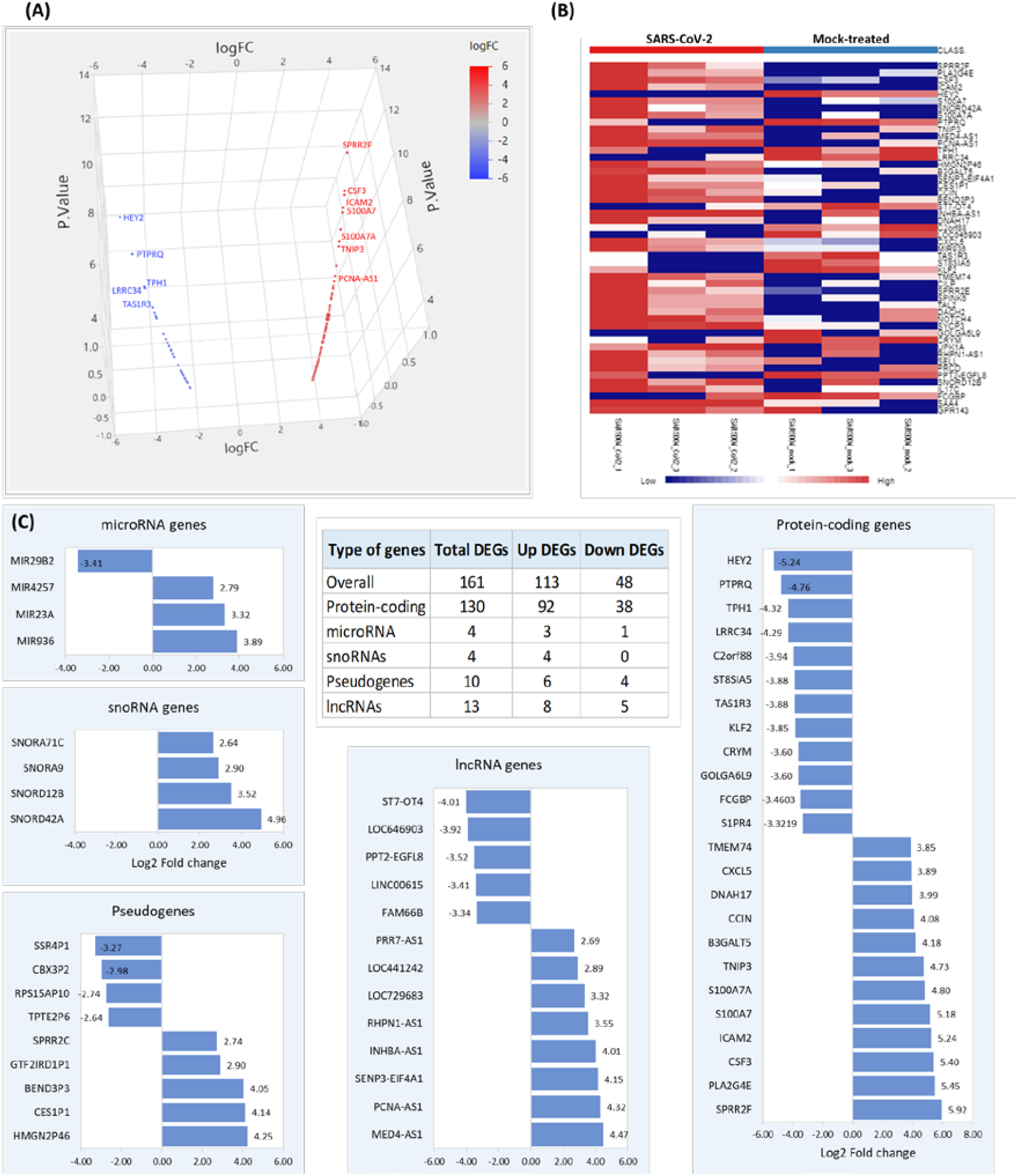
Functional annotations of differential expression genes (DEGs). (A) 3D-scatter plot for the upregulated (red) and downregulated (blue) DEGs. (B) Gene set enrichment analysis showing the top 100 variable genes between the six samples. (C) The numbers of the upregulated and downregulated differentially expressed genes. Relative expression level of DEGs stratified by the type of genes.

Of the down-regulated genes, (a) hes related family bHLH transcription factor with YRPW motif 2 (*HEY2*) is a nuclear transcription factor that represses DNA and negatively regulates miR-146a, IL-6, IL-1β, and TNF-α expression, (b) Protein Tyrosine Phosphatase Receptor Type Q (*PTPRQ*) phosphatase which catalyzes the dephosphorylation of phosphotyrosine and phosphatidylinositol and plays roles in cellular proliferation and differentiation and is required for auditory function, and (c) Taste 1 Receptor Member 3 (*TAS1R3*) encoding for G-protein coupled receptor involved in taste responses. The detailed information on DEGs is listed in **Supplementary Table S2**.

#### Functional annotations of genes

In order to analyze the aberrant gene expression pattern in SARS-CoV-2 infection, further functional analysis and annotation for DEGs were performed (**Figure 1C)**. Thirteen lncRNAs were deregulated following SARS-CoV-2 treatment. Annotation analysis revealed that *PPT2-EGFL8* readthrough (*PPT2-EGFL8*, FC=-3.5169|*p*=5.86E-06) was previously associated with circulating phospho- and sphingolipid concentrations, some other were associated with gastric, breast, and prostate cancer as INHBA-antisense RNA 1 (*INHBA-AS1*, FC=4.0|*p*=2.41E-07), RHPN1 antisense RNA 1 (*RHPN1-AS1*, FC= 3.54|*p*=4.85E-06), and ST7 overlapping transcript 4 (*ST7-OT*, FC=4-4.01|*p*=2.35E-07).

Genes for four miRNAs were deregulated; miR-936 (FC=3.88|*p*=5.4E-07), miR-23a (FC=3.32|*p*=1.8E-05), and miR-4257 (FC=2.79, *p*=3.1E-04) were upregulated, while miR-29b-2 was downregulated (FC=-3.4|*p*=1.12E-05). Four small nucleolar RNA genes were upregulated; including two small nucleolar RNA, C/D box: *SNORD42A* (FC=4.95|*p*=1.69E-10) and *SNORD12B* (FC=3.51|*p*=5.86E-06), and two small nucleolar RNA, H/ACA box: *SNORA9* (FC=2.90|*p*=1.8E-04) and *SNORA71C* (FC=2.63|*p*=6.8E-04). In addition, ten pseudogenes (6 upregulated and 4 downregulated DEGs) were identified to be deregulated. Despite being non-functional DNA segments that resemble functional genes, some might contain inherited or acquired promoter elements and exert beneficial regulatory function (www.GeneCards.org).

DEGs included 130 protein-coding genes (92 up and 38 down). Annotation of these genes revealed to enclose 7 transcription factors; three were activated; (a) TAL bHLH transcription factor 2 (*TAL2*), a basic helix-loop-helix transcription factor (FC=3.74|*p*=4.7E-04), (b) dachshund family transcription factor 2 (*DACH2*), a winged helix/forkhead transcription factor (FC=3.68|*p*=0.0006), and (c) AF4/FMR2 family member 2 (*AFF2*), a DNA-binding transcription factor (FC=3.40|*p*=0.0023), and four were downregulated, (a) *HEY2*, basic helix-loop-helix transcription factor (FC=-5.24|*p*=3.44E-08), and three C2H2 zinc finger transcription factor, (b) Kruppel like factor 2 (*KLF2*, FC=-3.85|*p*=2.6E-04), (c) early growth response 2 (*EGR2*, FC=-2.83|*p*=0.024), and (d) zinc finger and BTB domain containing 16 (*ZBTB16*, FC=-2.74|*p*=0.0327). Furthermore, three genes for extracellular matrix structural protein were under-expressed; (a) IgGFc-binding protein (*FCGBP*, FD=-3.46, *p*=8.26E-06), (b) Collagen alpha-3(VI) chain (*COL6A3*, FC=-2.93|*p*=0.00015), and (c) Complement component C1q receptor (*CD93*, FC=-2.74, *p*=0.0004). Additionally, phospholipid-transporting ATPase IM (*ATP8B4*), an active transporter (FC=-2.88|*p*=0.0002) and Retroviral-like aspartic protease 1 (*ASPRV1*), a viral or transposable element protein, (FC=-2.83|*p*=0.0002) were also downregulated. Two serine protease inhibitors were also downregulated; Insulin-like growth factor-binding protein 5 (*IGFBP5*, FC=-2.65|*p*=0.0006) and Serpin B10 (*SERPINB10*, FC=-2.81|*p*=0.00028).

In contrast, multiple proteases were upregulated in SARS-CoV-2 infection compared to mock-treated samples. Of these, SENP3-EIF4A1 readthrough gene (*SENP3-EIF4A1*, FC=4.15|*p*=8.93E-08), Chymotrypsin-like protease (*CTRL-1*, FC=2.88|*p*=0.00019), T-cell differentiation antigen *CD6* (FC=2.88, *p*=0.00021), *HTRA4* (FC=2.92, *p*=0.00016), *ABHD1* (FC=2.83|*p*=0.0002), and Cytosolic carboxypeptidase 3 *AGBL3* (FC=2.85|*p*=0.00024). Moreover, of the upregulated DEGs, the active transporter, Aquaporin (*AQP7*, FC=3.07, *p*=0.0095), the gap junction protein gamma 2 (*GJC2*, FC=3.32|*p*=1.87E-05), multiple membrane traffic proteins as MX dynamin like GTPase 1 (*MX1*, FC=2.71|*p*=0.0004) and synaptotagmin 5 (*SYT5*, FC=2.88|*p*=0.0002), and ion channels as gamma-aminobutyric acid type A receptor subunit rho2 (*GABRR2*, FC=2.74|*p*=0.0004) and bestrophin 4 (*BEST4*, FC=2.91|*p*=0.00017) (**Supplementary Table S2**).

#### Functional enrichment analysis of DEGs

To explore the functions of the DEGs, the 113 upregulated and 48 downregulated genes were subjected to GO and KEGG pathway enrichment analysis. As shown in **Figure 2A-C**, the significantly enriched GO terms for upregulated genes were mainly related to the acute inflammatory response (GO:0002526|FDR=6.46E-08), response to virus (GO:0009615|FDR=7.27E-08), regulation of immune response (GO:0050776|FDR=2.23E-08), negative regulation of viral genome replication (GO:0045071|FDR=1.60E-07), in addition to cytokine activity (GO:0005125|FDR=2.66E-09), chemokine activity (GO:0008009|FDR=0.0018), chemotaxis (GO:0006935|FDR=5.07E-07), and response to stress (GO:0006950|FDR=8.37E-14) (**Supplementary Table S3**). KEGG pathway analysis (**Figure 2D**) showed that upregulated genes were mainly enriched in IL-17 signaling pathway (KEGG ID: hsa04657|FDR=1.39E-15), TNF signaling pathway (hsa04668|FDR=5.34E-15), NOD-like receptor signaling pathway (hsa04621|FDR=4.07E-10), Cytokine-cytokine receptor interaction (hsa04060|FDR=2.08E-07), Jak-STAT signaling pathway (hsa04630|FDR=5.38E-06), NF-κB signaling pathway (hsa04064|FDR=1.33E-07), and Influenza A (hsa05164|FDR=1.10E-07). Other viral pathways included Kaposi’s sarcoma-associated herpesvirus infection (hsa05167), Herpes simplex infection (hsa05168), Measles (hsa05162), Hepatitis C (hsa05160), and Epstein-Barr virus infection (hsa05169) (**Supplementary Table S4**). In contrast, no significant GO terms or pathways were significant for the downregulated genes.

**Fig. 2.**
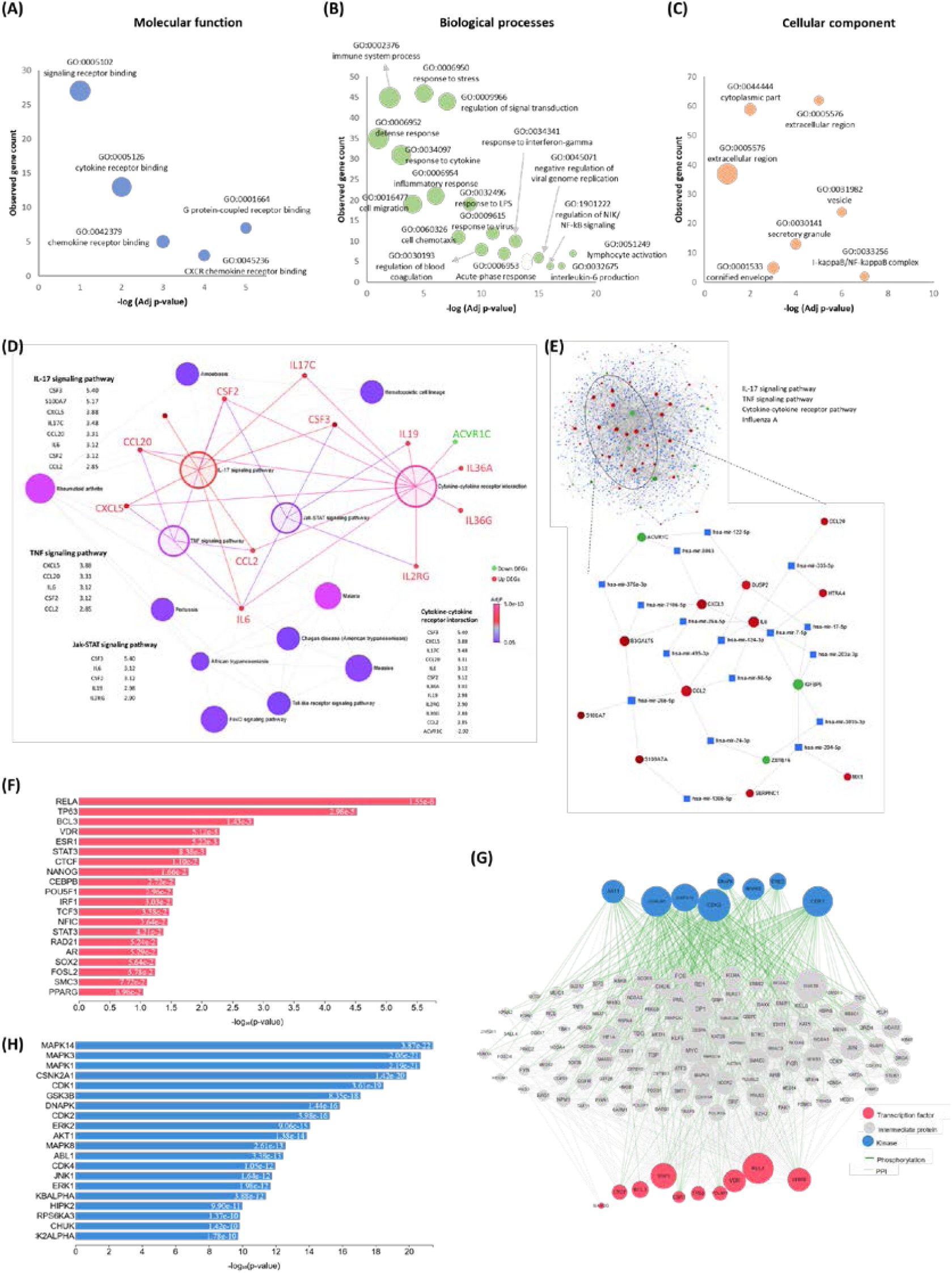
Functional enrichment analysis and gene regulatory networks. (A) Gene ontology analysis for molecular function, (B) for biological processes, (C) for cellular components, (D) KEGG Pathway enrichment analysis. Nodes represented pathways, with color based on its significance. Four enlarged highlighted nodes represented the most significant pathways. Bipartite gene networks with these nodes showed the association with upregulated (red) and downregulated (green) genes. (E) Gene-microRNAs interaction. Data source for interaction pairs: TarBase and miRTarBase. Red and green circles represented upregulated and downregulated DEGs respectively. Blue squares represented the microRNAs. Full network is shown with extracted nodes (ellipse) for four significant KEGG pathways. (F) Transcription Factor Enrichment Analysis (TFEA) showing putative transcription factors that most likely to regulate the differences in gene expression. (G) Upstream regulatory network that connects the enriched transcription factors to kinases through known protein-protein interactions. (H) Kinase Enrichment Analysis (KEA). Candidate enriched protein kinases that most likely regulate the formation of the identified transcriptional complexes. They are ranked based on overlap between known kinase–substrate phosphorylation interactions and the proteins in the protein-protein interaction subnetwork created in Figure 2G.

As depicted in **Figure 2D**, upregulated pathways are highly connected with inflammatory cytokines exhibiting significant crosstalk in particular. *IL-6* (also known as B-stimulatory factor-2) is involved in the final differentiation of B cells into immunoglobulin-secreting cells and the induction of acute-phase reactants. *IL-6* has sequence similarity with granulocyte colony-stimulating factors 2 and 3 (*CSF2* and *CSF3*), which are also highly expressed following SARS-CoV-2 infection. Further, several interleukins (*IL17C, IL19, IL36A*, and *IL36G*) noted serve as proinflammatory cytokines for the regulation of dendritic cells and T cells, as well as Interleukin-2 Receptor Subunit Gamma (*IL2RG*), which encodes a common gamma chain essential for IL-receptor function. Overexpression of 2 members of the CC chemokine family were observed: C-C Motif Chemokine Ligand 2 (*CCL2*) and C-C Motif Chemokine Ligand 2 (*CCL20*), which function to induce the migration of monocytes and lymphocytes, respectively (www.GeneCards.org)

#### Gene regulatory networks

Gene-miRNA interactions network was constructed (**Figure 2E**). It consisted of 77 seeds (significant genes), 1136 nodes, and 1776 edges mapped to the corresponding molecular interaction databases. KEGG pathway enrichment analysis of the network was significant for four pathways; namely IL-17 signaling pathway (*p*=7E-05, Hits=5/93), TNF signaling pathway (*p*=0.0017, Hits=4/110), cytokine-cytokine receptor pathway (*p*=0.012, Hits=5/294), and Influenza A (*p*=0.044, Hits=3/167). After extraction of nodes relevant to these immune-related pathways, the densely connected microRNAs in the cluster module included: miR-26b-5p, miR-26a-5p, miR-124-3p, miR-7-5p, miR-17-5p, miR-335-5p, mirR-24-3p, miR-203a-3p, and miR-122-5p.

Upstream regulatory cell signaling layers responsible for the observed pattern in gene expression following SARS-CoV-2 infection is depicted in **Figure 2F-H**. Network included integrated transcription factors and kinases with PPI. The top enriched transcription factors were *RELA* proto-oncogene (*p*=1.55E-06), *BCL3* (*p*=0.0014), *TP63* (*p*=2.9E-05) and V*DR* (*p*=0.0051). *RELA*, a NF-KB Subunit, has a key role in mediating inflammation, differentiation, cell growth, tumorigenesis and apoptosis (www.GeneCards.org). It is associated with 12 overlapping targets in the DEGs list (*IER3, STAT1, NFKB2, ICAM1, DRAM1, NEDD9, IL32, NFKBIA, TNFAIP3, NFKBIZ, TNIP1*). Another NF-kB regulator, *BCL3*, which has paradox effect according to its subcellular localization; regulates transcriptional activation of NF-kappa-B target gene while being in the nucleus and inhibits the nuclear translocation of the NF-kappa-B p50 subunit when existed in the cytoplasm (www.GeneCards.org). From the DEGs defined following SARS-CoV-2 infection, six genes (*IER3, STAT1, NFKB2, NFKBIA, IRF9*, and *BIRC3*) were targeted by *BCL3*. The nuclear receptor for calcitriol, *VDR*, was also significantly associated with the up-regulated DEGs (*NFKB2, NFKBIA*, and *IRAK2*).

#### Tuberculosis pathway

Gene set enrichment analysis (GSEA) showed over-representation of SARS-CoV-2 genes in Tuberculosis pathway (ID: hsa05152, *p*=6.09E-04). In this particular pathway, 116 DEGS (of 179 genes) were upregulated (**Supplementary Figure S2)**. *IL6, TLR9*, and *TNF* were at the top of that list. *VDR* and *CEPB* are further noted for their critical regulation of inflammatory and immune responses, including SARS-CoV-2 infected cells.

### Transcriptomic changes following BCG vaccine administration

#### Differential expressed genes

After preprocessing of the expression data from the four different stages post vaccination (**Supplementary Figure S3**), 696, 66, 49, and 80 DEGs were detected at 14 days, 28 days, 56 days, and 84 days respectively. As depicted in **Figure 3**, principal component analysis showed clustering with clear demarcation from the initial stage at day zero. A total of 696, 66, 49, and 80 DEGs respectively were identified (**Figure 3D**). The altered expression patterns across 4 different time points showed host immune responses to be constantly changing post-BCG vaccination. Stratified analysis according to the direction of DEGs showed similar results with no overlapping between up and down DEGs at each stage (**Figure 3E-F**).

**Fig. 3.**
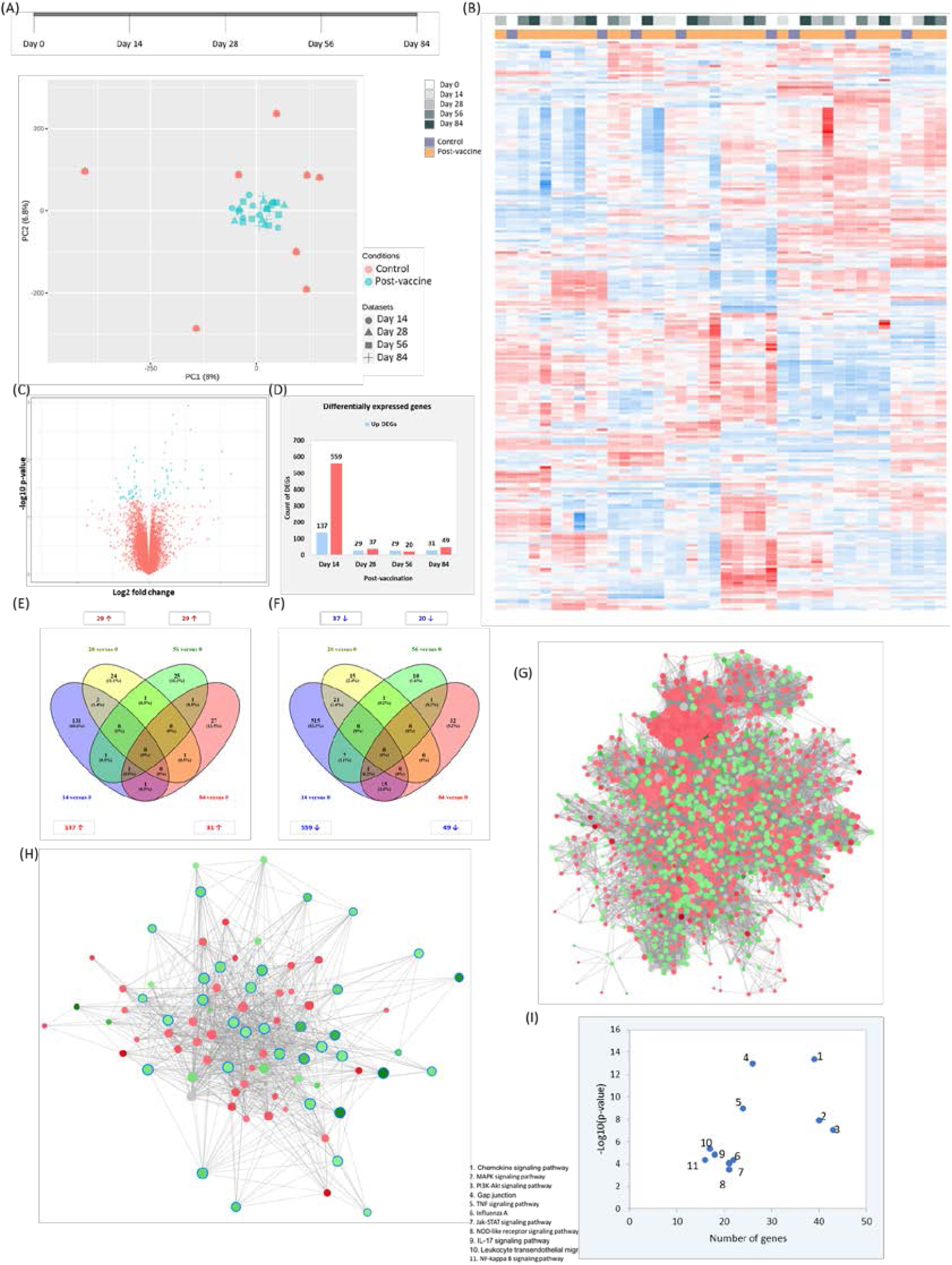
Differential expression genes and functional enrichment analysis following BCG vaccination. (A) Principal component analysis after normalization, showing cluster of samples post-vaccination on day 14 through day 84. (B) Clustergram showing hierarchical clustering of the top 2500 variable genes among the five groups. (C) Volcano plot representing log 2-fold change and − log 10 (adjusted P-value). (D) Number of DEGs at each stage post-vaccination (day 14 versus day 0, day 28 versus day 0, day 56 versus day 0, and day 84 versus day 0) (E-F) Venn diagram showing intersection between upregulated and downregulated DEGs at different stage. Number of upregulated and downregulated DEGs are demonstrated in blue and red boxes, respectively. (G) Protein-protein interaction network for DEGs following BCG vaccination. String interactome for PPI showing up-regulated (red nodes) and down-regulated (green nodes) genes. (H) A cluster of inhibited KEGG pathways. Circled in blue the gene list of the selected pathways. (I) Top significant pathways inhibited following BCG vaccination (enriched in the cluster).

#### Functional enrichment analysis

GO enrichment analysis was carried out in three categories, including biological processes (BP), molecular functions (MF), and cellular components (CC). The most significant terms for BP, MF, and CC were the translation (GO:0006412|FDR=8.59E-10), RNA binding (GO:0003723|FDR=6.98E-08), and Cytosolic ribosome (GO:0022626|FDR=1.30E-08) at Day 14; short-chain fatty acid catabolic process (GO:0019626|FDR=0.0008), azole transmembrane transporter activity (GO:1901474|FDR=0.0088), and nucleolar ribonuclease P complex (GO:0005655|FDR=0.1409) at Day 28; protein import into peroxisome matrix (GO:0016558|FDR=0.0029), beta-1,3-galactosyltransferase activity (GO:0048531|FDR=0.0046), and manchette (GO:0002177|FDR=0.0121) at Day 56; and positive regulation of T-helper 2 cell differentiation (GO:0045630|FDR=0.0088), primary lysosome (GO:0005766|FDR=0.0246), and alanine transmembrane transporter activity (GO:0022858|FDR=0.0121) at Day 84 post vaccination.

Pathway enrichment analysis showed multiple activated pathways at different stages. The most significant upregulated pathways at an early stage (Day 14 and Day 28 groups) were (a) Ribosome (hsa03010, FDR=3.83E-12) including multiple mitochondrial ribosomal proteins (as *MRPL17, MRPS21, RPL23A, MRPL24, MRPL34, MRPL36*), and ribosomal proteins for small subunits (*RPS14, RPS15, RPS16, RPS19, RPS28*), and large subunits (*RPL18A, RPL36, RPLP2, RPL13, RPL15*) and (b) Propionyl-CoA catabolism (R-HSA-71032|FDR= 0.0001) via modulating methylmalonyl CoA epimerase (*MCEE*), Methylmalonic aciduria type A (*MMAA*), and propionyl-CoA carboxylase (*PCCB*) genes. However, at later stages, the top upregulated pathways were Aminoacyl-tRNA biosynthesis (hsa00970|FDR=0.0246), Scavenging of heme from plasma (R-HSA-2168880|FDR=0.0056), and Heme Biosynthesis (WP561|FDR=0.020).

In contrast, of the inhibited pathways after 2-weeks of vaccination were Cytokine-cytokine receptor interaction (hsa04060|FDR=2.63E-04), JAK-STAT signaling pathway (hsa04630|FDR=0.047), and ECM-receptor interaction (hsa04512|FDR=0.05456). Cell adhesion molecules (hsa04514|FDR=0.0055) and Cytokines and Inflammatory Response (WP530|FDR=0.137) were downregulated in the same cohorts one month after vaccination. However, delayed host immune response at Day 56 showed downregulation of NOD-like receptor signaling pathway (hsa04621|FDR=0.0029), PI3K-Akt signaling pathway (hsa04151|FDR=0.0057), and Signaling by Interleukins (R-HSA-449147|FDR=0.0048), while inhibited Interferon Signaling (R-HSA-913531|FDR=0.0014) and Interferon alpha/beta signaling (R-HSA-909733|FDR=0.0015) were inactivated in patients at Day 84 post vaccination. Activated and inhibited GO and pathway terms for each stage are listed in detail in **Supplementary Tables S5-S12.**

#### Protein-protein interaction network

On comparing combined post-vaccination samples (days 14, 28, 56, and 84) versus day 0, 76 DEGs (41 up and 35 down) were defined. Protein-protein interaction network was constructed from the combined DEGs. The clusters with densely connected nodes in the PPI network were detected (**Figure 3G**). KEGG enrichment analysis revealed 121 significant downregulated pathways. Eleven pathways were extracted in a separate subnetwork module; namely Influenza A, IL-17 signaling pathway, TNF signaling pathway, Chemokine signaling pathway, NOD-like receptor signaling pathway, PI3K-Akt signaling pathway, NF-κB signaling pathway, JAK-STAT signaling pathway, MAPK signaling pathway, Gap junction, and Leukocyte transendothelial migration (**Figure 3H-I**). The number of nodes was 2076, while corresponding edges counted for 6123. To identify hub genes involved in the host immune response following BCG vaccination, further gene set enrichment analysis of the cluster was performed. Downregulated genes with high strength of enrichment as *NFKB1* (Degree of connectivity with other genes=91), *RELA* (Degree=95), *IKBKB* (Degree=60) genes were enriched in 8 (out of 11) pathways, followed by *PIK3CA* (Degree=119), PIK3CD (Degree=59), *PRKCB* (Degree=26), *RAF1* (Degree=69), *AKT1* (Degree=221) and *AKT2* (Degree=48) were involved in 6 (out of 11) pathways.

### Comparison between transcriptomic signature in SARS-CoV-2 infection and BCG vaccination

#### Intersecting common pathways

A total of 52 enriched KEGG pathways were upregulated in SARS-CoV-2 infection. Cross-reference of these upregulated pathways with downregulated KEGG pathways in BCG vaccination experiment, 45 common pathways were intersected accounting for 86.5% of SARS-CoV-2 upregulated pathways (**Figure 4A-C**). These pathways were categorized into the following groups: (1) cellular processes, (2) organismal systems, (3) environmental information, and (4) human diseases. The top significant ones in SARS-CoV-2 experiment were IL-17 signaling pathway (FDR=1.39E-15|Hits=14/92; overlapping genes in the pathway compared to the total gene set in the pathway), TNF signaling pathway (FDR=5.34E-15|Hits=14/108 genes), NOD-like receptor signaling pathway (FDR=4.07E-10|Hits=12/166 genes). Other immune-related pathways were Cytokine-cytokine receptor interaction (FDR=2.08E-07|Hits=11/263 genes), NF-κB signaling pathway (FDR=1.33E-07|Hits=8/93 genes), and JAK-STAT signaling pathway (FDR=5.38E-06|Hits=8/160 genes). In the same 45 pathways, viral infectious agents included Influenza A (FDR=1.10E-07|Hits=10/168 genes), Measles (FDR=1.33E-07|Hits=9/133 genes), Herpes simplex virus 1 infection (FDR=1.33E-07|Hits=10/181 genes) and bacterial infections included Legionellosis (FDR=0.0014|Hits=7/54 genes) and Tuberculosis (FDR=0.0039|Hits=5/172 genes). Overlapping genes within each pathway that were significantly upregulated in SARS-CoV-2 infection and downregulated following BCG vaccination at the four different time points (14, 28, 56, and 84 days), and are demonstrated in **Supplementary Table S13**. Although different genes were modified at each time point, the pathways remained constantly downregulated post-BCG vaccination.

**Fig. 4.**
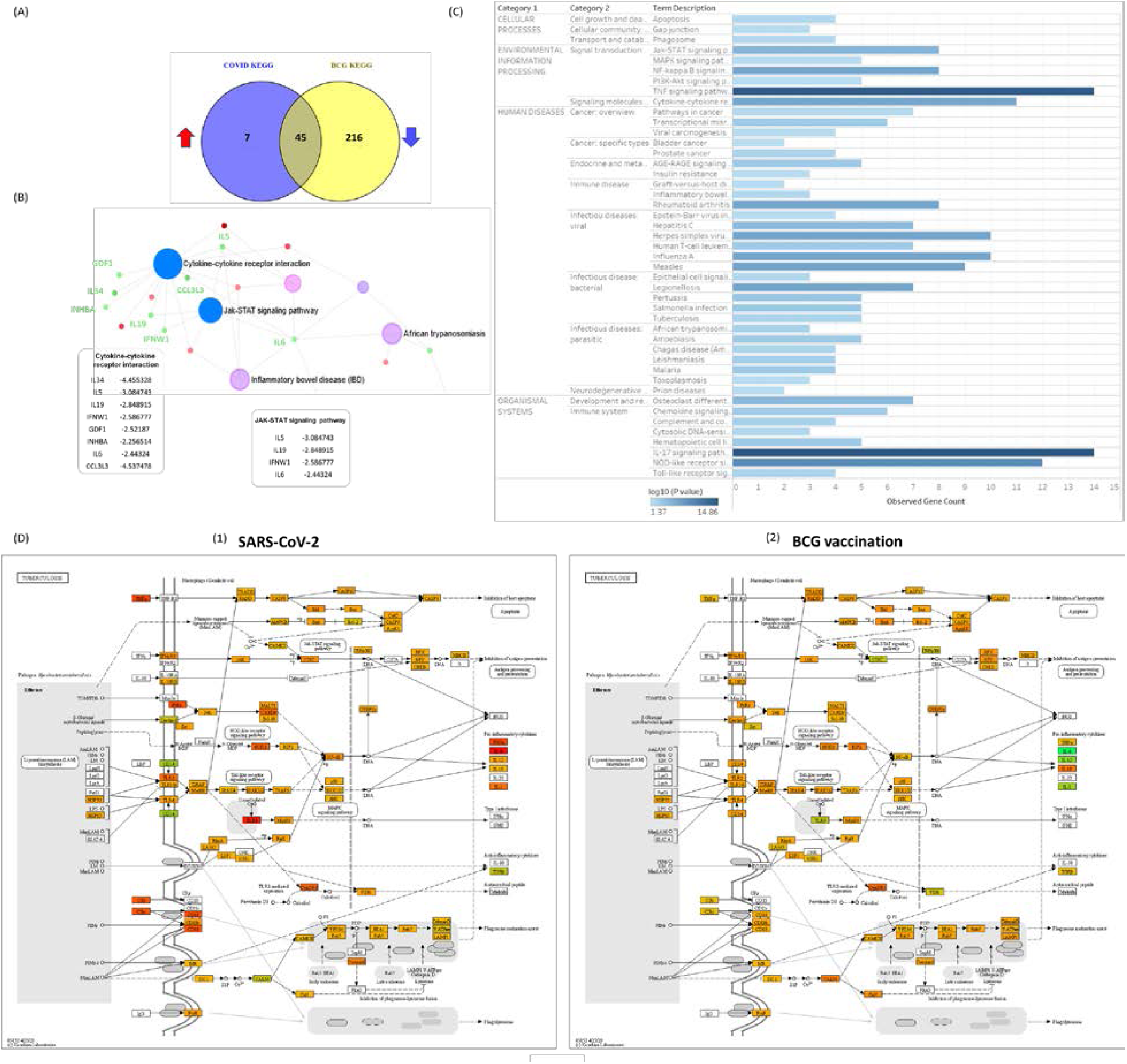
Common pathways between SARS-CoV-2 and BCG vaccination experiments. (A) Intersection between KEGG pathways of SARS-CoV-2 infected cells and BCG vaccination. (B) Expression level of some hub genes following BCG vaccination showing downregulation post-vaccination as an example for reversed direction of expression. (C) Displayed 45 upregulated KEGG signaling pathways in SARS-CoV-2 cell line which are downregulated following BCG vaccination. Bars represented the observed gene count for each pathway. Degree of color represents the degree of significance (the more intensity, the higher significance). KEGG signaling pathways are categorized according to the functional hierarchical classification system. (D) Expression intensity of genes in the Tuberculosis KEGG pathway. Colored by the log fold change of DEGs in (1) SARS-CoV-2 infection compared to mock-treated cells and (2) following BCG vaccination.

#### Comparative enrichment analysis in Tuberculosis pathway

On comparing 116 DEGs in SARS-CoV-2 infected cells which were over-represented in the 179 gene set of Tuberculosis pathways with their expression pattern after BCG vaccination, the top upregulated inflammatory markers; including *IL6/IL12, TLR1/2/9, TNF, C3, IL1A/B, NFKB1, TGFB2, AKT1*, and *HLA-DPA1/B1*, and transcription factors as *NFKB1* and *STAT1* showed reversed direction post-vaccination. Additionally, the down-regulated DEGs in SARS-CoV-2, including several non-receptor serine/threonine-protein kinase (*AKT3, CAMK2D*, and *MAPK10*), the monocyte differentiation antigen (*CD14*), the membrane trafficking regulatory protein (*LAMP2*), growth factor (*TGFB3*), the apoptotic regulator (*BCL2*), and proinflammatory cytokines (*IL18*) switched their direction following BCG vaccine administration (**Figure 4D**). Similar paradox directions of expression level in some DEGs for SARS-CoV-2 cells and BCG vaccine samples were also noted in **Supplementary Figure S4-S10**.

## DISCUSSION

### Overview

To gain insight into the engendering mechanism of the SARS-CoV-2 infection, the gene expression profiles of the virus were systematically analyzed through bioinformatic techniques. In this study, a total of 161 DEGs, including 113 upregulated and 48 downregulated, were screened. The biological functions of these DEGs were explored based on GO function and pathway enrichment data. Further analysis indicated significant upregulation in 45 distinct KEGG pathways in SARS-CoV-2 infection overlapped with pathways down-regulated following the BCG vaccination. In addition to the pathway analysis, this study describes potential kinases, transcription factors, miRNAs, and lncRNAs differentially expressed in Sars-CoV-2 infection (**Figure 2 and 5**). Several candidate DEGs, including *IL-6, CCL20, CSF2, ICAM*, and *CXCL1*/*2* were highlighted for their key roles in respiratory viral infection, tuberculosis, and overall immune function. To elucidate the relationship between COVID-19 infection and BCG-vaccination, the common pathways were categorized into the following groups: inflammatory and immunoregulatory, signaling, and infectious pathways.

### Common Inflammatory and Immunoregulatory Pathways

Of SARS-CoV-2 infected cells, analysis implicates several upregulated genes enriched in inflammatory pathways, most significantly in the IL-17 and NOD-like receptor pathways. Conversely, these same pathways were downregulated in the BCG-vaccinated group, suggesting BCG vaccination may compensate for aspects of pathway dysregulation in SARS-CoV-2 infection. The family of IL-17 mediates protective innate immunity against external pathogens and plays a central role in the self-clearance of intracellular pathogens^37,38^. Additionally, T helper cells (Th17), which themselves produce Il-17, are known to be key players in the pathogenesis of chronic inflammatory diseases and autoimmune tissue destruction^39^.

Elevated Th17 responses and IL-17 pathways are seen in COVID-19 patients and have been linked to ‘cytokine storms’, a surge of pro-inflammatory molecules that are associated with the Acute Respiratory Distress Syndrome (ARDS) typically seen in these patients^40^. BCG vaccination has been shown to confer protective immunity in the Rhesus macaque model by decreasing Th17 cell maturation, thus suppressing Th17 response and preventing production of IL-17^23,41^. Further, IL-17A and IL-17F induce proinflammatory gene expression of NF*-κ*B, MAPK, in synergy with TNF-α and IL-6. *IL-17* gene family also eliminates self-reactive T cells through production of granulocyte colony-stimulating factor (*G-CSF*) and several chemokines upregulated in Sars-CoV-2 infected cells, including *CXCL1, CXCL2, CCL20* and *IL-17*^39,42^ **(Figure 9**).

The other major upregulated inflammatory pathway, NOD-Like receptor signaling pathway, detects various pathogens and stimulates innate immune responses against them, driving the subsequent activation of cytokine, NF-*κ*B, and MAPK pathways. Upregulation of NOD-like receptor signaling without appropriate negative feedback regulation can contribute to pathological tissue damage^43^. Innate and adaptive immune response to an invading pathogen relies on the ability of the body to recognize foreign elements and is stimulated by inflammasomes, which are intracellular multi-protein complexes such as NOD-like receptors (NLR)^44^. *NLRC5* (short for NOD-like receptor family CARD domain containing 5) regulates major histocompatibility complex class I expression in the course of an infection. In accordance with this, a study reported knockdown of *NLRC5* resulted in decreased levels of CD8+ cells in influenza A virus^45^. Our study identifies significant upregulation of IL-17 and NOD-like receptor signaling pathways in the immune response of COVID-19, pathways that were markedly downregulated in BCG-vaccination.

### Common Signal Transduction Pathways

Similarly, several signal transduction pathways were up-regulated in Sars-CoV-2 and inversely downregulated in the BCG-vaccinated group with TNF, NF-κB, MAPK and JAK/STAT signaling pathways conferring the most statistical significance. TNF signaling upregulation is heavily involved in the complex regulation of immune cells. Following trimerization with either TNFR1 (expressed nearly ubiquitously) or TNFR2 (expressed mostly in immune cells), the TNF signaling pathway can activate NF-κB and MAPK signaling pathways downstream of it. Upon viral infection, NF-κB proteins regulate the transcription of many genes, including antimicrobial peptides, cytokines, chemokines, stress-response proteins and anti-apoptotic proteins^46^. In a study of Middle Eastern Respiratory Syndrome (MERS), a viral infection caused by a member of the coronavirus family, researchers found the virus downregulated antiviral cytokines (TNF-alpha) as it induced pro-inflammatory cytokines (Il-1 beta, Il-6, Il-8) during initial infection^47^. In a study of Severe Acute Respiratory Syndrome (SARS), yet another member of the coronavirus family, TNF-alpha was similarly downregulated and induced NF-κB activation^46^. In the SARS pathogen, ACE2 was determined to be the functional receptor of the virus through the regulation of TNF-alpha, which ultimately permitted viral entry and promoted respiratory pathogenesis. Altogether, the literature on other members of the coronavirus family indicates a disruption of TNF-signaling pathway, which is in accordance with the marked pathway upregulation of TNF pathways signaling noted in our findings. In BCG vaccinated models *in vitro*, exposure of human peripheral blood mononuclear cells (PBMCs) to BCG treatment boosted IL-6 and TNF-α expression in response to lipopolysaccharide stimulation^22^. Similarly, BCG-immunized adults produced high TNF-α and IL-1β expression 3-month post-vaccination^23^. In addition to sustained cytokine response at 3 months, the production of heterologous Th1 and Th17 remained elevated 1-year post-BCG vaccination, concluding the vaccine created long-term immune responses to pathogens besides *M.tb* alone^23^.

Downstream of TNF, the NF-κB signaling pathway similarly affects a broad range of biological processes, including adaptive immune, inflammatory, and stress responses. Upon viral infection, NF-κB proteins regulate the transcription of many genes, including antimicrobial peptides, cytokines, chemokines, stress-response proteins and anti-apoptotic proteins^46^. Of the COVID-19 upregulated genes involved in the NF-κB pathway were *CSF-2* and *IL-6*, transcription factors and innate immune system mediators^55,56^. The upregulation of *CSF*-*2*, produced by endothelial and immune cells, is similarly upregulated in other respiratory diseases, including *mycoplasma pneumoniae* and *M.tb*, by promoting neutrophil and macrophage inflammatory response. The upregulation of IL-6 has been noted in other members of the coronavirus family, including the pathogenic agent responsible for SARS. In murine models, SARS-coronavirus spike protein, which is evolutionarily conserved in Sars-CoV-2, induced upregulation of IL-6 through the NF-κB pathway. Higher levels of proinflammatory cytokines, including IL-6, TNF, CSF-2, IL-1b, and IL-8 are noted in COVID-19 patients with increasing levels as a predictor of the severity of pneumonia^57^.

Additionally, MAPK signaling was upregulated in the Sars-CoV-2 infected group while inversely downregulated in the BCG-vaccinated group. MAPK regulates cellular processes of proliferation, stress responses, as well as immune defense^44,48,49^. MAPK signaling cascade is activated in response to external stress signals of the 3 MAP kinases (ERK, JNK, and p38 isoforms) with the subsequent stimulation of pro-inflammatory cytokines by JNK and p38 signaling^50-52^. In the nucleus, activated JNK and p38 promote multiple effector proteins, including NF-κB, c-Jun, STAT1^53^. These effector proteins have been implicated in viral infections such as influenza A and HSV-1 and their dysregulation by pathogens is associated with impaired antiviral response by the host^53,54^. The downregulation of the MAPK pathway in the BCG-vaccinated group (**Figure 10**) may be able to partially reverse the upregulation induced by the SARS-CoV-2 virus. Additionally, our results indicated the top 3 upregulated kinases in SARS-CoV-2 group were *MAKP14, MAPK3*, and *MAPK1*, further corroborating the central role MAPK plays in viral pathogenesis of SARS-CoV-2 infection (**Figure 6**).

JAK-STAT signaling, which is a major pathway involved in cytokine signaling and subsequent inflammation, was upregulated in SARS-CoV-2 infection as well. IFNs induce the JAK/STAT signaling pathway, which in turn activates transcription of widespread immune protection. For this reason, several viruses have evolved to target the JAK/STAT pathway themselves^58^. Several studies have recently advocated for the potential benefit of commercially available Jak 1 and 2 inhibitors as anti-inflammatory treatment in COVID-19 cases^58^; however, there is concern from the scientific community that JAK inhibitors may promote the evolution of SARS-CoV-2 virus as this has been reported in herpes viruses. BCG vaccination may be key in attenuating the JAK/STAT pathway without entirely impairing interferon-mediated response. *In vitro* experiments show that BCG vaccination induces 2 members of the suppressor of cytokine signaling (SOCS) family, SOCS1/3, eliciting a negative feedback regulator of the JAK/STAT signaling cascade via IFN-gamma regulation^59^.

Overall, although TNF, NF-κB, MAPK, and JAK/STAT signaling were upregulated in SARS-CoV-2 infection, they were markedly downregulated in BCG, suggesting BCG vaccination may be targeting these same pathways. The exact mechanism remains elusive, but identification of these pathways and their targets are key to mitigating pathogenesis and merit further investigation.

### Infectious and Human Disease Signaling Pathways

Further, in the current study, BCG vaccination down regulated several pathways related to viral and bacterial inflammatory disease pathways including influenza A, measles, Herpes simplex virus 1, Legionellosis, and tuberculosis, that were up-regulated in SARS-CoV-2 infection. Prior studies have demonstrated non-specific effects of BCG against viral infection^21^, and bacterial infections^32, 60^. In one study, mice inoculated with BCG displayed overall increased resistance to encephalomyocarditis, murine hepatitis, type 1 and 2 herpes simplex, foot-and-mouth disease and A0 and A2 influenza viruses^61^. Intercellular adhesion molecules (or CD54), included in the upregulated DEGs, are receptors utilized for pathogen entry into host cells that promote intercellular signaling and survival of virus in the host cell. In human rhinovirus infection and influenza virus, ICAM-1 is involved in viral protein uncoating, delivering the viral RNA genome into the cytoplasm of host cells across a lipid bilayer^62,63^. Other studies corroborate this showing that most viruses interact with and induce ICAM-1 expression, including HIV, Human Parainfluenza virus, and rhinovirus infection^64-66^. Presence of cytokines such as IFN-γ, TGF-beta, and TNF-α induces expression of ICAM on macrophage for activation of T cells, this was observed in human TB infected macrophages^67^. BCG immunotherapy of bladder tumors cells showed mycobacteria infection-induced expression of ICAM-1 molecules, thereby eliciting immune response through improved antigen presentation to T lymphocytes^68^. BCG is thought to inhibit inflammasome activation via Zinc metalloproteases in order to improve immunogenicity^60,69^ and remains the current standard therapy for non-muscle invasive bladder cancer.

### The Role of VDR and Other Enriched Transcription Factors

Compared to mock-treated cells, SARS-CoV-2 infected cells exhibited significant enrichment of *VDR* transcription factor, one of the major downstream signaling transduction nuclear receptor in tuberculosis KEGG pathway, and was significantly associated with the up-regulated DEGs, namely *NFKB2, NFKBIA*, and *IRAK2*, which are related to NF-κB and Toll-like receptor signaling pathways. Normally, VDR binds to calcitriol, the active form of Vitamin D. The VDR-calcitriol complex interacts with retinoid-X-receptor to form a heterodimer capable of regulating transcriptional activity, leading to several pleiotropic transcriptional effects. Similarly, *VDR* gene expression was continuously upregulated during HIV^70^ and influenza virus infection^71^. Previously reported, VDR signaling may repress cytokine gene expression in activated T-cells, consequently reducing inflammatory response^28,72^. Part of the mechanism of action of BCG vaccine against tuberculosis is through increased production of Vitamin D, which binds with its nuclear receptor *VDR*, resulting in the generation of antimicrobial peptides (cathelicidin and beta-defensin) and death of intracellular *M.tb*^44^. In short, Vitamin D plays an important role in modulating the innate and adaptive immune response alongside its classically characterized role in bone health. Long before the scientific basis was understood, Vitamin D was unknowingly and empirically being used to help tuberculosis patients who were sent to sanatoriums for sun-light exposure and prescribed cod-liver - remedies rich in Vitamin D^73^. Since then, Vitamin D has been well characterized for its immuno-modulatory properties, including the regulation of IFN-gamma, a key activator of macrophage response^74^ and promoting the production of regulatory T cells^75^. In addition to its role in *M.tb*^76^, cathelicidin is a highly conserved protein that has been shown to direct antiviral activity in several respiratory viral infections including respiratory syncytial virus, human Rhinovirus (HRV), and influenza A^77-82^. *In vitro*, Vitamin D has been shown to counteract the *M.tb*-induced downregulation of cathelicidin by actually recovering cathelicidin levels and promoting Th1 cell differentiation. Following BCG vaccination, Vitamin D levels remain elevated, indicating Vitamin D is being upregulated long after the initial inoculation. It has been postulated that increased calcitriol recruits dendritic cells from the site of inoculation to the lymph nodes, accounting for the sustained response^24^. This serves as a possible mechanism of protective effects of BCG vaccination in relation to consistently elevated plasma Vitamin D concentration present for as long as 9 months as reported by the authors^24^. A major complication of COVID-19 infection is inflammation of the air sacs of the lungs resulting in pneumonia. Vitamin D deficiency contributed to the pathogenesis of acute respiratory distress syndrome (ARDS)^83^. Biotransformation into the active form of vitamin D-1,25-dihydroxyvitamin D (calcitriol) by the 1-α-hydroxylase (CYP27B1) enzyme found in liver cells, and immune cells expression of *VDR*, confirms major role of Vitamin D in immune regulation; both in innate and adaptive immune responses^25-28,72.^. Mutation in the *VDR* gene single nucleotide polymorphism reported in children was associated with response to viral infection in children^82^. Given these findings, the transcriptional upregulation of *VDR* in SARS-CoV-9 may be contributing to the dysregulation of this response by upregulating cytokine response, interrupting microbial peptides, and promoting immune invasion in the lungs; therefore, the well characterized role of *VDR* in BCG vaccination could serve as a potential biomarker against COVID-19 disease.

In addition to *VDR*, our bioinformatic analysis showed deregulation of multiple transcription factors in SARS-CoV-2 infection. Specifically, *TAL2, DACH2*, and *AFF2* were upregulated whose functional role in viral immune response is unclear, while *HEY2, KLF2, EGR2*, and *ZBTB16* were under-expressed and were previously reported to have eminent role in fine-tuning immune response. *HEY2* was the most downregulated gene in our list. It is a known transcription repressor, modulating cardiovascular development, neurogenesis, and oncogenesis^84^. A recent study demonstrated the role of *HEY2* in inflammation, namely chronic periodontitis, via negative regulation of IL-6, IL-1β, and TNF-α expressions^85^. *KLF2* is a key player in T cell differentiation, trafficking, quiescence, and survival, in addition to the regulation of endothelial function. Mice with *KLF2* deficiency experienced an activated T cell phenotype with severely reduced T cells in the periphery and increased susceptibility to HIV-1 infection^86,87^. *EGR2* transcription factor can control adaptive innate immunity and lead to functional impairment of T cells^88^. After infection with influenza virus, *EGR2* knockout mice exhibited prolonged viral shedding, impaired CD4 + T-cell response, infiltration of memory precursor type CD8 + T cells into the lung with decreased IFN-γ, TNFα, and granzyme B^89^. *ZBTB16* transcription factor can mediate the ubiquitination and subsequent proteasomal degradation of target proteins^90^. Transcription factor enrichment analysis unravels multiple transcription factors modulating the deregulated genes in SARS-CoV-2 infected cells. *RELA*, a well-known antiviral transcription factor, was the top enriched transcription factor for DEGs following SARS-CoV-2 infection. It is previously reported to promote the growth of cytopathic RNA viruses by extending the lifespan of infected cells and serve as the replicative niche of intracellular pathogens^91^. *TP63* was also shown to be a cellular regulator of the HPV life cycle^92^.

### MicroRNAs, small nucleolar RNAs and pseudogenes

In the present study, we observed some deregulated miRNAs and highlighted the relationship between DEGs and miRNAs networking which may account for the transcriptomic changes evident in SARS-CoV-2 expression. Of particular note, mir-26a-5p, miR-26b-5p, and miR-124-3p, which have all been well characterized for their involvement in viral and bacterial inflammatory pathways, particularly Influenza A, RSV, and *M.tb*, were noted here for their putative respiratory pathogenesis. In *M.tb* infection, miR-26a-5p can modulate macrophage IFN-gamma responsiveness^3^. In another study, this same microRNA was shown to be downregulated in Influenza A virus in humans, contributing to viral pathogenesis^4^. The other family member, miR-26b-5p, can inhibit viral replication in Vesicular stomatitis virus (VSV) and Sendai virus (SeV) by inducing type-1 IFN expression^5^. RSV acts to upregulate miR-26b, which in turn inhibits Toll-like receptor TLR4, a key sentinel in adaptive immune response ^6^. A similar effect on TLRs is seen in the pathogenesis of *M.tb* in alveolar macrophages wherein microRNA-124 negatively regulates TLR signaling. SARS-CoV-2 infected cells exhibited overexpression of miR-23a, which was previously shown to promote the replication of human herpes simplex virus type 1 via downregulating interferon regulatory factor 1 (IRF1), an innate antiviral molecule^97^. Altogether, the highlighted cluster of microRNAs are likely regulating viral pathways relevant to SARS-CoV-2 infection. Post-transcriptional regulation of some genes altered in SARS-CoV-2 infected cells are therefore worthy of experimental validation.

Small nucleolar RNAs and pseudogenes were found to be deregulated in SARS-CoV-2 cells. SnoRNAs can serve as a source of short regulatory RNA species involved in the control of processing and translation of various mRNAs. Despite their exact role in COVID-19 and tuberculosis is unclear and complicated, these ncRNAs were found to be upregulated in virus-infected human cells. They can guide chemical modifications of structural RNAs and have a role in cell-cell communication^98^. SnoRNAs were reported to have a paradox effect; they can act as mediators of host antiviral response, and on the other hand, are utilized by viruses to evade innate immunity and complete their life cycle^98^. Furthermore, different groups of pseudogenes were activated in human cells upon triggering inflammation with known cellular protein, including virus particles and chunks of bacterial cell walls, highlighting the unique functions of pseudogenes in host immune response^99^.

### Limitations

To the best of our knowledge, our study is the first to dig into the regulatory mechanisms underlying the host immune response against SARS-CoV-2infection and to provide a putative protective role of BCG vaccination. Altogether, several DEGs with a high connectivity degree in the PPI network were identified in this study, which showed the strong inverse correlation in SARS-CoV-2 infection and BCG vaccination with upregulation in the former and downregulation in the latter. This inverse relationship suggests BCG vaccination may mediate key pathways in SARS-CoV-2 infection through NSEs. Nevertheless, it is essential to recognize some limitations of administering BCG vaccines, including, the heterogeneity of BCG response due to pharmacogenomic variation of patients, contraindications of BCG-vaccination for immunosuppressed patients, and the presence of different strains of BCG vaccines in the market^100^. Although the exact mechanisms with which BCG elicits protective effects are not entirely understood, its efficacy in infectious, oncogenic, and inflammatory disease are well documented and may confer similar effects if COVID-19.

## Conclusion

In summary, bioinformatic approach identified key differentially expressed genes involved in the pathogenesis of COVID-19. Significant nodes in the PPI network, including high degrees in IL-17 signaling, TNF signaling pathway, NOD-like receptor, and NF-κB give novel insight into the inflammatory and signal transduction pathways mediating SARS-CoV-2 viral infection. The 45 common pathways upregulated in SARS-CoV-19 and downregulated in BCG vaccination suggest BCG vaccination may help to mitigate this pathway dysregulation. Although further experimental validation is warranted to verify our discoveries, the results give insight into potential biomarkers and targeted therapy to address the ongoing COVID-19 pandemic. We sustain BCG vaccination may be a safe and cost-effective alternative in incurring partial protection against COVID-19 pandemic until more targeted measures can be produced and implemented.

## Contributors

ET and EK designed the study. ET and MH analyzed the data. ET and EK interpreted the data. ET, JS, and TD search literature and wrote the manuscript. All authors reviewed and approved the final version of the manuscript.

## Supplementary Materials

Supplementary Materials (https://drive.google.com/open?id=15Na738L282XNaQAJUh0cZf1WoG9jJfzJ)

### Supplementary Figures

Supplementary Fig. S1. Quality check of HNBE cell lines. (A) Density plot against log2 of read counts to display the relative distribution of different counts in each group. (B) Box plot showing distribution of raw read counts after normalization. (C) Principal Component Analysis to demonstrate the distribution pattern of infected and mock treated samples. (D) Volcano plot showing distribution of genes according to their fold change and significance.

Supplementary Fig. S2. Gene set enrichment analysis in Tuberculosis pathway (KEGG ID: hsa05152). (A) Expression intensity of genes in the Tuberculosis KEGG pathway in three SARS-CoV-2 infected cells and mock-treated cells. (B) Enrichment of differentially expressed genes (blue) in Tuberculosis pathway. (C) Fold change of up-regulated TB-related genes following SARS-CoV-2 infection compared to mock-treated cell lines.

Supplementary Fig. S3. Data exploration and quality check after normalization of RNA seq data (GSE87186). (A) Box plot for the 40 samples included in the dataset. (B) Principal component analysis showing variations in the data. Each point represents a sample. (C) Density plot showing uniform distribution between samples.

Supplementary Figure S4. Expression level of DEGs in Interleukin 17 signaling pathway. Colored by the log fold change of DEGs (A) DEGs in SARS-CoV-2 infection compared to mock-treated cells, (B) DEGs following BCG vaccination.

Supplementary Figure S5. Expression level of DEGs in TNF signaling pathway. Colored by the log fold change of DEGs (A) DEGs in SARS-CoV-2 infection compared to mock-treated cells, (B) DEGs following BCG vaccination.

Supplementary Figure S6. Expression level of DEGs in NOD-like receptor signaling pathway. Colored by the log fold change of DEGs (A) DEGs in SARS-CoV-2 infection compared to mock-treated cells, (B) DEGs following BCG vaccination.

Supplementary Figure S7. Expression level of DEGs in cytokine-cytokine receptor interaction pathway. Colored by the log fold change of DEGs (A) DEGs in SARS-CoV-2 infection compared to mock-treated cells, (B) DEGs following BCG vaccination.

Supplementary Figure 8. Expression level of DEGs in JAK-STAT signaling pathway. Colored by the log fold change of DEGs (A) DEGs in SARS-CoV-2 infection compared to mock-treated cells, (B) DEGs following BCG vaccination.

Supplementary Figure S9. Expression level of DEGs in NF-Kappa B signaling pathway. Colored by the log fold change of DEGs (A) DEGs in SARS-CoV-2 infection compared to mock-treated cells, (B) DEGs following BCG vaccination.

Supplementary Figure S10. Expression level of DEGs in Influenza A pathway. Colored by the log fold change of DEGs (A) DEGs in SARS-CoV-2 infection compared to mock-treated cells, (B) DEGs following BCG vaccination.

### Supplementary Tables

Supplementary Table S1. Raw expression data in SARS-CoV-2 infected human normal bronchial epithelial cell line compared to mock-treated cells.

Supplementary Table S2. Differentially expressed genes in SARS-CoV-2 infected human normal bronchial epithelial cell line compared to mock-treated cells.

Supplementary Table S3. Significant Gene Ontology terms in SARS-CoV-2 infected cells compared to mock-treated cells.

Supplementary Table S4. Functional enrichment pathway analysis in SARS-CoV-2 infected cells compared to mock-treated cells.

Supplementary Table S5. Top 10 enriched gene ontology functions of the differentially expressed genes at day 14 following BCG vaccination.

Supplementary Table S6. Top 10 enriched pathways of the differentially expressed genes at day 14 following BCG vaccination.

Supplementary Table S7. Top 10 enriched gene ontology functions of the differentially expressed genes at day 28 following BCG vaccination.

Supplementary Table S8. Top 10 enriched pathways of the differentially expressed genes at day 28 following BCG vaccination.

Supplementary Table S9. Top 10 enriched gene ontology functions of the differentially expressed genes at day 56 following BCG vaccination.

Supplementary Table S10. Top 10 enriched pathways of the differentially expressed genes at day 56 following BCG vaccination.

Supplementary Table S11. Top 10 enriched gene ontology functions of the differentially expressed genes at day 84 following BCG vaccination.

Supplementary Table S12. Top 10 enriched pathways of the differentially expressed genes at day 84 following BCG vaccination.

Supplementary Table S13. Intersected upregulated KEGG pathways in SARS-COV-2 infected cells that are reversed following BCG vaccination.

## REFERENCES

1. Kissler, S. M., Tedijanto, C., Goldstein, E., Grad, Y. & Lipsitch, M. Projecting the transmission dynamics of SARS-CoV-2 through the postpandemic period. Science 5793, 1–13 (2020).

2. Chinazzi M. et al., The effect of travel restrictions on the spread of the 2019 novel coronavirus (COVID-19) outbreak. Science 10.1126/science.aba9757 (2020).

3. Kucharski, A. J. et al. Early dynamics of transmission and control of COVID-19: a mathematical modelling study. Lancet Infect. Dis. https://doi.org/10.1016/S1473-3099(20)30144-4 (2020)

4. Harbert, R., Cunningham, S. W. & Tessler, M. Spatial modeling cannot currently differentiate SARS-CoV-2 coronavirus and human distributions on the basis of climate in the United States. (2020).

5. Deshwal, V. K. COVID 19: A comparative study of Asian, European, American continent. Int. J. Sci. Res. Eng. Dev. 3, 436–440 (2020)

6. Jiang, S., Hillyer, C. & Du, L. Neutralizing antibodies against SARS-CoV-2 and other human coronaviruses. Trends Immunol., 1–5 (2020). doi: 10.1016/j.it.2020.03.007

7. Walls, A. C. et al. Structure, function, and antigenicity of the SARS-CoV-2 spike glycoprotein. Cell 180, 1–12 (2020).

8. Hoffmann, M. et al. SARS-CoV-2 cell entry depends on ACE2 and TMPRSS2 and is blocked by a clinically proven protease inhibitor. Cell 181, 1–10 (2020).

9. Li, W. et al. Angiotensin-converting enzyme 2 is a functional receptor for the SARS coronavirus. Nature 426, 450–454 (2003).

10. Wu, F. et al. A new coronavirus associated with human respiratory disease in China. Nature 579, 265–269. doi: 10.1038/s41586-020-2008-3

11. Wang, Q. et al. Structural and functional basis of SARS-CoV-2 entry by using human ACE2. Cell doi: 10.1016/j.cell.2020.03.045 (2020)

12. Ong Z. E. et al. A dynamic immune response shapes COVID-19 progression. Cell Press doi: 10.1016/j.chom.2020.03.021 (2020)

13. Zhang, C., Wu, Z., Li, J., Zhao, H. & Wang, G. The cytokine release syndrome (CRS) of severe COVID-19 and Interleukin-6 receptor (IL-6R) antagonist Tocilizumab may be the key to reduce the mortality. Int J Antimicrob. Agents 105954 doi: 10.1016/j.ijantimicag.2020.105954 (2020)

14. Hegarty, P. K., Service, N. H., Kamat, A. M. & Dinardo, A. BCG vaccination may be protective against Covid-19. medRxiv doi: 10.13140/RG.2.2.35948.10880 (2020)

15. Gursel, M. & Gursel, I. Is Global BCG vaccination coverage relevant to the progression of SARS-CoV-2 pandemic? Medical Hypotheses doi: https://doi.org/10.1016/j.mehy.2020.109707 (2020)

16. Dayal, D. & Gupta, S. Connecting BCG vaccination and COVID-19: Additional Data. medRxiv 2755657, 2020.04.07.20053272 (2020).

17. Miller, A., Reandelar, M. J., Fasciglione, K., Roumenova, V., Li, Y. & Otazu, G. H. Correlation between universal BCG vaccination policy and reduced morbidity and mortality for COVID-19: an epidemiological study. J Chem. Inform. Model. 53, 1689–1699 (2012)

18. Shet, A., Ray, D., Malavige, N., Santosham, M. & Bar-Zeev, N. Differential COVID-19-attributable mortality and BCG vaccine use in countries. medRxiv doi: 10.1101/2020.04.01.20049478 (2020)

19. Berg, M. K., Yu, Q., Salvador, C. E., Melani, I., Kitayama, S. Mandated Bacillus Calmette-Guérin (BCG) vaccination predicts flattened curves for the spread of COVID-19. medRxiv doi: https://doi.org/10.1101/2020.04.05.20054163 (2020)

20. Shann, F. Nonspecific effects of vaccines and the reduction of mortality in children. Clinical Ther. 35, 109–114 (2013). https://doi.org/10.1016/j.clinthera.2013.01.007

21. Moorlag, S. J., Arts, R. J., van Crevel, R., Netea, M. G. Non-specific effects of BCG vaccine on viral infections. Clin. Microbiol. Infect. 25, 1473–1478 (2019).

22. Netea, M. G., Schlitzer, A., Placek, K., Joosten, L. A. & Schultze, J. L. Innate and Adaptive Immune Memory: an Evolutionary Continuum in the Host’s Response to Pathogens. Cell Host Microbe 25, 13–26 (2019)

23. Kleinnijenhuis, J. et al. Long-lasting effects of BCG vaccination on both heterologous Th1/Th17 responses and innate trained immunity. J. Innate Immun. 6, 152–158 (2014).

24. Lalor, M. K. et al. Population differences in immune responses to Bacille Calmette-Guérin vaccination in infancy. J. Infect. Dis. 199, 795–800 (2009). doi: 10.1086/597069

25. McMahon, L. et al. Vitamin D-mediated induction of innate immunity in gingival epithelial cells. Infect. Imm. 79, 2250–2256 (2011).

26. Lagishetty, V., Liu, N. Q. & Hewison, M. Vitamin D metabolism and innate immunity. Mol. Cell Endocrinol. 347, 97–105 (2011).

27. Prietl, B., Treiber, G., Pieber, T. R. & Amrein, K. Vitamin D and immune function. Nutrients 5, 2502–2521 (2013).

28. Lin, R. Crosstalk between vitamin D metabolism, VDR signalling, and innate immunity. BioMed. Res. Int. Article ID 1375858, 5 pages (2016). http://dx.doi.org/10.1155/2016/1375858

29. Hensel, J., Mcandrews, K. M., Mcgrail, D. J. & Dowlatshahi, D. P. Exercising caution in correlating COVID-19 incidence and mortality rates with BCG vaccination policies due to variable rates of SARS CoV-2 testing. medRxiv https://doi.org/10.1101/2020.04.08.20056051 (2020)

30. Anthony, S. J. et al. Global patterns in coronavirus diversity. Virus Evol. 3, 1–15 (2017).

31. Kirov, S. Association between BCG policy is significantly confounded by age and is unlikely to alter infection or mortality rates. medRxiv https://doi.org/10.1101/2020.04.06.20055616 (2020)

32. Arts, R. J. W. et al. BCG vaccination protects against experimental viral infection in humans through the induction of cytokines associated with trained immunity. Cell Host Microbe 23, 89-100.e5 (2018).

33. Lachmann, A. et al. Massive mining of publicly available RNA-seq data from human and mouse. Nature Comm 9, 1366 (2018)

34. Ashburner, M. et al. Gene ontology: tool for the unification of biology. Nat. Genet. 25, 25e9 (2000).

35. Kanehisa, M. & Goto. S. Kegg: Kyoto encyclopedia of genes and genomes. Nucleic Acids Res. 28, 27e30 (2000).

36. Chen, E. Y. et al. Enrichr: interactive and collaborative HTML5 gene list enrichment analysis tool. BMC Bioinform 128, (2013).

37. Cua, D. J. & Tato, C. M. Innate IL-17-producing cells: the sentinels of the immune system. Nat. Rev. Immunol. 10, 479–488 (2010).

38. Rudner, X. L., Happel, K. I., Young, E. A. & Shellito, J. E. Interleukin-23 (IL-23)-IL-17 cytokine axis in murine *Pneumocystis carinii* infection. Infect. Immun. 75, 3055–3061 (2007).

39. Kuwabara, T., Ishikawa, F., Kondo, M. & Kakiuchi, T. The Role of IL-17 and related cytokines in inflammatory autoimmune diseases. Mediators Inflamm. doi: 10.1155/2017/3908061 (2017)

40. Wu, D. & Yang, X. TH17 responses in cytokine storm of COVID-19: An emerging target of JAK2 inhibitor Fedratinib. J. Microb. Immun. Infect. doi: 10.1016/j.jmii.2020.03.005 (2020)

41. Dijkman, K. et al. Prevention of tuberculosis infection and disease by local BCG in repeatedly exposed rhesus macaques. Nat. Med. 25, 255–262 (2019).

42. Ye, P. et al. Requirement of interleukin 17 receptor signaling for lung CXC chemokine and granulocyte colony-stimulating factor expression, neutrophil recruitment, and host defense. J. Exp. Med. 194, 519–527 (2001).

43. Zhong, Z. et al. NF-κB restricts inflammasome activation via elimination of damaged mitochondria. Cell 164, 896–910 (2016).

44. Liu, T., Zhang, L., Joo, D. & Sun, S. C. NF-κB signaling in inflammation. Signal Transduct. Target Ther. 2, 17023 (2017).

45. Lupfer, C. R., Stokes, K. L., Kuriakose, T. & Kanneganti, T. Deficiency of the NOD-Like receptor NLRC5 results in decreased CD8+ T cell function and impaired viral clearance. J. Virol. 91, e00377–17 (2017).

46. Wang, W., et al. Up-regulation of IL-6 and TNF-α induced by SARS-coronavirus spike protein in murine macrophages via NF-κB pathway. Virus Res. 128, 1–8 (2007).

47. Kindrachuk, J. et al. Antiviral potential of ERK/MAPK and PI3K/AKT/mTOR signaling modulation for Middle East Respiratory Syndrome coronavirus infection as identified by temporal kinome analysis. Antimicrob. Agents Chemother. 59, 1088–1099 (2015).

48. Dong, C., Davis, R. J. & Flavell, R. A. MAP kinases in the immune response. Ann. Rev. Immunol. 20, 55–72 (2002).

49. Arthur, J. S. C., & Ley, S. C. Mitogen-activated protein kinases in innate immunity. Nat. Rev. Immunol. 13, 679–692 (2013).

50. Kaminska, B. MAPK signalling pathways as molecular targets for anti-inflammatory therapy--from molecular mechanisms to therapeutic benefits. Biochim. Biophys. Acta 1754, 253–262 (2005).

51. Zhang, Y., & Dong, C. Regulatory mechanisms of mitogen-activated kinase signaling. Cell. Mol. Life Sci. 64, 2771–2789 (2007).

52. Soares-Silva, M., Diniz, F. F., Gomes, G. N. & Bahia D. The mitogen-activated protein kinase (MAPK) pathway: role in immune evasion by Trypanosomatids. Frontiers Microbiol. 7, 183 (2016).

53. Kumar, R. et al. Role of MAPK/MNK1 signaling in virus replication. Virus Res. 253, 48–61 (2020).

54. Hirasawa, K., Kim, A., Han, H. S., Han, J., Jun, H. S. & Yoon, J. W. Effect of p38 mitogen-activated protein kinase on the replication of encephalomyocarditis virus. J. Virol. 77, 5649–5656 (2003).

55. Begum, N. A. et al. Mycobacterium bovis BCG cell wall-specific differentially expressed genes identified by differential display and cDNA subtraction in human macrophages. Infect. Immun. 72, 937–948 (2004).

56. Verreck, F. A., de Boer, T., Langenberg, D. M., Zanden, L. & Ottenhoff, T. H. Phenotypic and functional profiling of human proinflammatory type-1 and anti-inflammatory type-2 macrophages in response to microbial antigens and IFN-gamma- and CD40L-mediated costimulation. J. Leukoc. Biol. 79, 285–293 (2006).

57. Chen, G. et al. Clinical and immunological features of severe and moderate coronavirus disease 2019. J. Clin. Invest. 1–10 (2020). https://doi.org/10.1172/JCI137244

58. Richardson, P. et al. Baricitinib as potential treatment for 2019-nCoV acute respiratory disease. Lancet 395, e30–e31 (2020). doi: 10.1016/S0140-6736(20)30304-4

59. Imai, K., Kurita-Ochiai, T. & Ochiai, K. *Mycobacterium bovis* bacillus Calmette-Gueérin infection promotes SOCS induction and inhibits IFN-γ-stimulated JAK/STAT signaling in J774 macrophages, FEMS Immunol. Med. Microb. 39, 173–180 (2003)

60. Iqbal, N. T. & Hussain, R. Non-specific immunity of BCG vaccine: A perspective of BCG immunotherapy. Trials Vaccinol. 3, 143–149 (2014).

61. Floc’h, F. & Werner, G. H. Increased resistance to virus infections of mice inoculated with BCG (Bacillus calmette-guérin). Ann. Immunol. 127, 173–186 (1976).

62. Othumpangat, S., Noti, J. D., McMillen, C. M. & Beezhold, D. H. ICAM-1 regulates the survival of influenza virus in lung epithelial cells during the early stages of infection. Virol. 487, 85–94 (2016).

63. Bella, J. & Rossmann, M. G. ICAM-1 receptors and cold viruses. Pharmacochem. Lib. 31, 291–297 (2000).

64. Christensen, J. P., Johansen, J., Marker, O. & Thomsen, A. R. Circulating intercellular adhesion molecule-1 (ICAM-1) as an early and sensitive marker for virus-induced T cell activation. Clin. Exp. Immunol. 102, 268–273 (1995).

65. Gao, J., Choudhary, S., Banerjee, A. K. & De, B. P. Human parainfluenza virus type 3 upregulates ICAM-1 (CD54) expression in a cytokine-independent manner. Gene Expr. 9, 115–121 (2000).

66. Scheglovitova, O. et al. Antibody to ICAM-1 mediates enhancement of HIV-1 infection of human endothelial cells. Arch. Virol. 140, 951–958 (1995).

67. DesJardin, L. E., Kaufman, T. M., Potts, B., Kutzbach, B., Yi, H. & Schlesinger, L. S. Mycobacterium tuberculosis-infected human macrophages exhibit enhanced cellular adhesion with increased expression of LFA-1 and ICAM-1 and reduced expression and/or function of complement receptors, FcgammaRII and the mannose receptor. Microbiol. 148, 3161–3171 (2002).

68. Jackson, A. M., Alexandroff, A. B., McIntyre, M., Esuvaranathan, K., James, K. & Chisholm, G. D. Induction of ICAM 1 expression on bladder tumours by BCG immunotherapy. J. Clin. Pathol. 47, 309–312 (1994).

69. Master, S. S. et al. Mycobacterium tuberculosis prevents inflammasome activation. Cell Host Microbe 3, 224–232 (2008).

70. Nevado, J., Tenbaum, S. P., Castillo, A. I., Sánchez-Pacheco, A. & Aranda, A. Activation of the human immunodeficiency virus type I long terminal repeat by 1 alpha, 25-dihydroxyvitamin D3. J. Mol. Endocrinol. 38, 587–601 (2007).

71. Rieder, F. J. J. et al. Human cytomegalovirus infection downregulates vitamin-D receptor in mammalian cells. J. Steroid Biochem. Mol. Biol. 165, 356–362 (2017).

72. White, J. H. Vitamin D signaling, infectious diseases, and regulation of innate immunity. Infect. Immun. 76, 3837–3843 (2008).

73. Jarrett, P. & Scragg, R. A short history of phototherapy, vitamin D and skin disease. Photochem. Photobiol. Sci. 16, 283–290 (2017).

74. Lemire, J. M., Archer, D. C., Beck, L. & Spiegelberg, H. L. Immunosuppressive actions of 1,25-dihydroxyvitamin D3: preferential inhibition of Th1 functions. J. Nutr. 125, 1704S–1708S (1995).

75. Griffin, M. D., Xing, N. & Kumar, R. Vitamin D and its analogs as regulators of immune activation and antigen presentation. Ann. Rev. Nutr. 23, 117–145 (2003).

76. Sonawane, A. et al. Cathelicidin is involved in the intracellular killing of mycobacteria in macrophages. Cell Microbiol. 13, 1601–1617 (2011).

77. Sousa, F. H. et al. Cathelicidins display conserved direct antiviral activity towards rhinovirus. Peptides 95, 76–83 (2017).

78. Currie, S. M. et al. The human cathelicidin LL-37 has antiviral activity against respiratory syncytial virus. PLoS One 8, e73659 (2013) doi: 10.1371/journal.pone.0073659

79. Gunville, C. F., Mourani, P. M. & Ginde, A. A. The role of vitamin D in prevention and treatment of infection. Inflamm. Allergy Drug Targets 12, 239–245 (2013).

80. Liu, P. T. et al. Toll-like receptor triggering of a vitamin D-mediated human antimicrobial response. Science 311, 1770–1773 (2006). doi: 10.1126/science.1123933.

81. Roth, D. E. et al. Association between vitamin D receptor gene polymorphisms and response to treatment of pulmonary tuberculosis. J. Infect. Dis. 190, 920–927 (2004).

82. Kresfelder, T. L. et al. Confirmation of an association between single nucleotide polymorphisms in the VDR gene with respiratory syncytial virus-related disease in South African children. J. Med. Virol. 83, 1834–1840 (2011).

83. Dancer, R. C. et al. Vitamin D deficiency contributes directly to the acute respiratory distress syndrome (ARDS). Thorax 70, 617–624 (2015).

84. Fischer, A. & Gessler, M. Hey genes in cardiovascular development. Trends Cardiov. Med. 13, 221–226 (2003).

85. Lina, S., Lihong, Q., Di, Y., Bo, Y., Xiaolin, L., Jing, M. microRNA-146a and Hey2 form a mutual negative feedback loop to regulate the inflammatory response in chronic apical periodontitis. J. Cell Biochem. 120, 645–657 (2018).

86. Pearson, R., Fleetwood, J., Eaton, S., Crossley, M. & Bao, S. Krüppel-like transcription factors: a functional family. Int J. Biochem. Cell Biol. 40, 1996–2001 (2008).

87. Carlson, C. M. et al. Kruppel-like factor 2 regulates thymocyte and T-cell migration. Nature 442, 299–302 (2006).

88. Miao, T. et al. Egr2 and 3 control adaptive immune responses by temporally uncoupling expansion from T cell differentiation. J. Exp. Med. 214, 1787–1808 (2017).

89. Du, N. et al. Role for EGR2 in the T-cell response to influenza. Proceed. Nat. Acad. Sci. 111, 16484–16489 (2014).

90. Furukawa, M., He, Y. J., Borchers, C. & Xiong, Y. Targeting of protein ubiquitination by BTB-Cullin 3-Roc1 ubiquitin ligases. Nat. Cell Biol. 5, 1001–1007 (2003).

91. Bais, S. S. et al. Chandipura Virus Utilizes the Prosurvival Function of RelA NF-κB for Its Propagation. J. Virol. 93, e00081–19. (2019). doi: 10.1128/JVI.00081-19

92. Mighty, K. K. & Laimins, L. A. p63 is necessary for the activation of human papillomavirus late viral functions upon epithelial differentiation. J. Virol. 85, 8863–8869 (2011).

93. Ni, B., Rajaram, M. V., Lafuse, W. P., Landes, M. B. & Schlesinger, L. S. Mycobacterium tuberculosis decreases human macrophage IFN-γ responsiveness through miR-132 and miR-26a. J. Immunol. 193, 4537–4547 (2014).

94. Tambyah, P. A. et al. microRNAs in circulation are altered in response to influenza A virus infection in humans. PLoS One 8, e76811 (2013) doi: 10.1371/journal.pone.0076811

95. Liu, C., Zhang, L., Xu, R. & Zheng, H. MiR-26b inhibits virus replication through positively regulating interferon signaling. Viral Immunol. 31, 676–682 (2018).

96. Liu, S., Gao, L., Wang, X. & Xing, Y. Respiratory syncytial virus infection inhibits TLR4 signaling via up-regulation of miR-26b. Cell Biol. Int. 39, 1376–1383 (2015).

97. Ru, J. et al. MiR-23a facilitates the replication of HSV-1 through the suppression of interferon regulatory factor 1. PLoS One 9, e114021 (2014). doi: 10.1371/journal.pone.0114021

98. Stepanov, G. A., Filippova, J. A., Komissarov, A. B., Kuligina, E. V., Richter, V. A. & Semenov, D. V. Regulatory role of small nucleolar RNAs in human diseases. Biomed. Res. Int. 2015, 206849 (2015).

99. Rapicavoli, N. et al. 2013. A mammalian pseudogene lncRNA at the interface of inflammation and anti-inflammatory therapeutics. eLife 2, e00762 (2013).

100. Rowland, R. & McShane, H. Tuberculosis vaccines in clinical trials. Expert Rev. Vaccines 10, 645–658 (2011).

