## Supplementary figures for "Hidden in plain sight: The effects of BCG vaccination in COVID-19 pandemic"

Supplementary Table S13. Intersected upregulated KEGG pathways in SARS-COV-2 infected cells that are reversed following BCG vaccination.

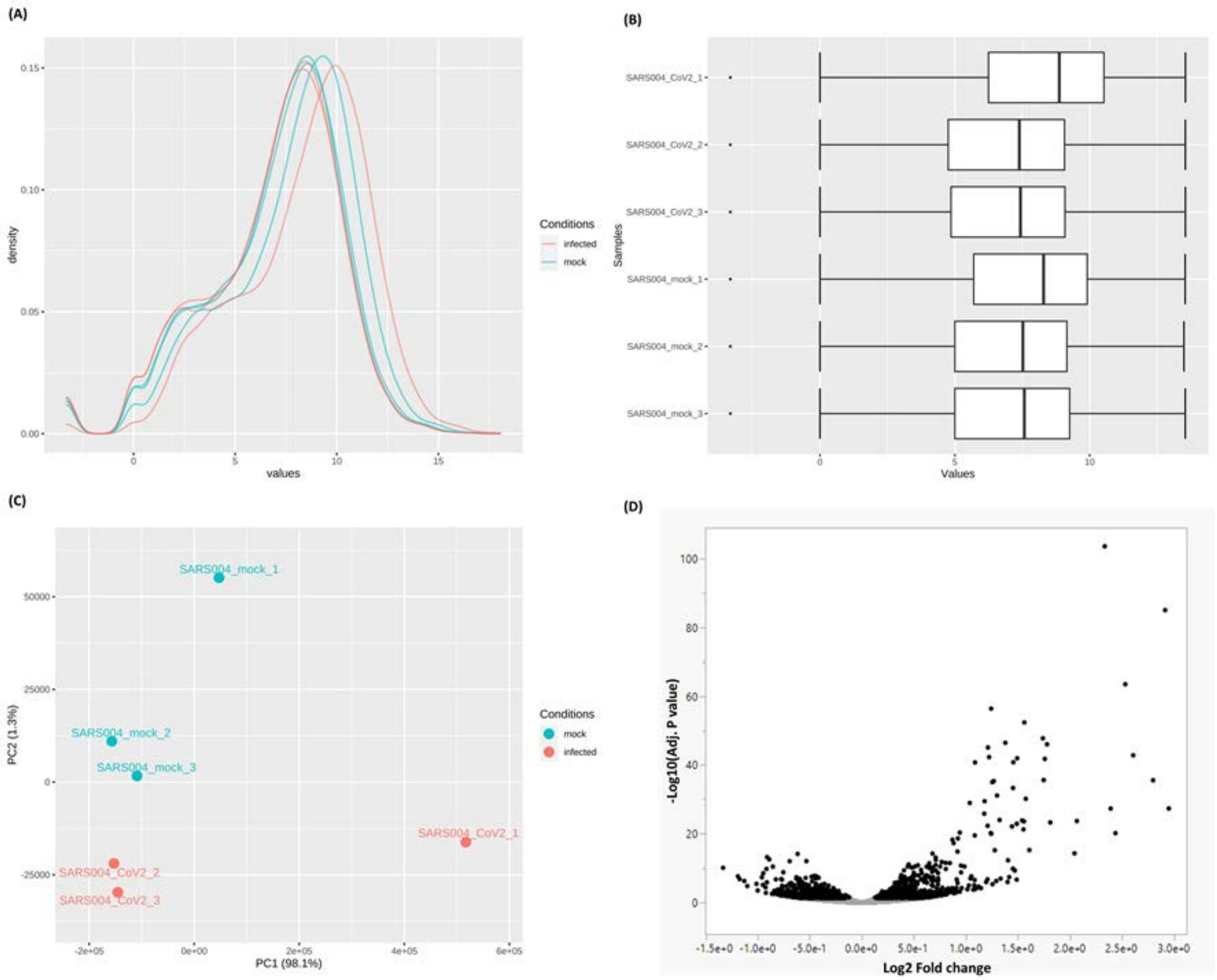


**Supplementary Fig. S1. Quality check of HNBE cell lines.** (A) Density plot against log2 of read counts to display the relative distribution of different counts in each group. (B) Box plot showing distribution of raw read counts after normalization. (C) Principal Component Analysis to demonstrate the distribution pattern of infected and mock treated samples. (D) Volcano plot showing distribution of genes according to their fold change and significance.

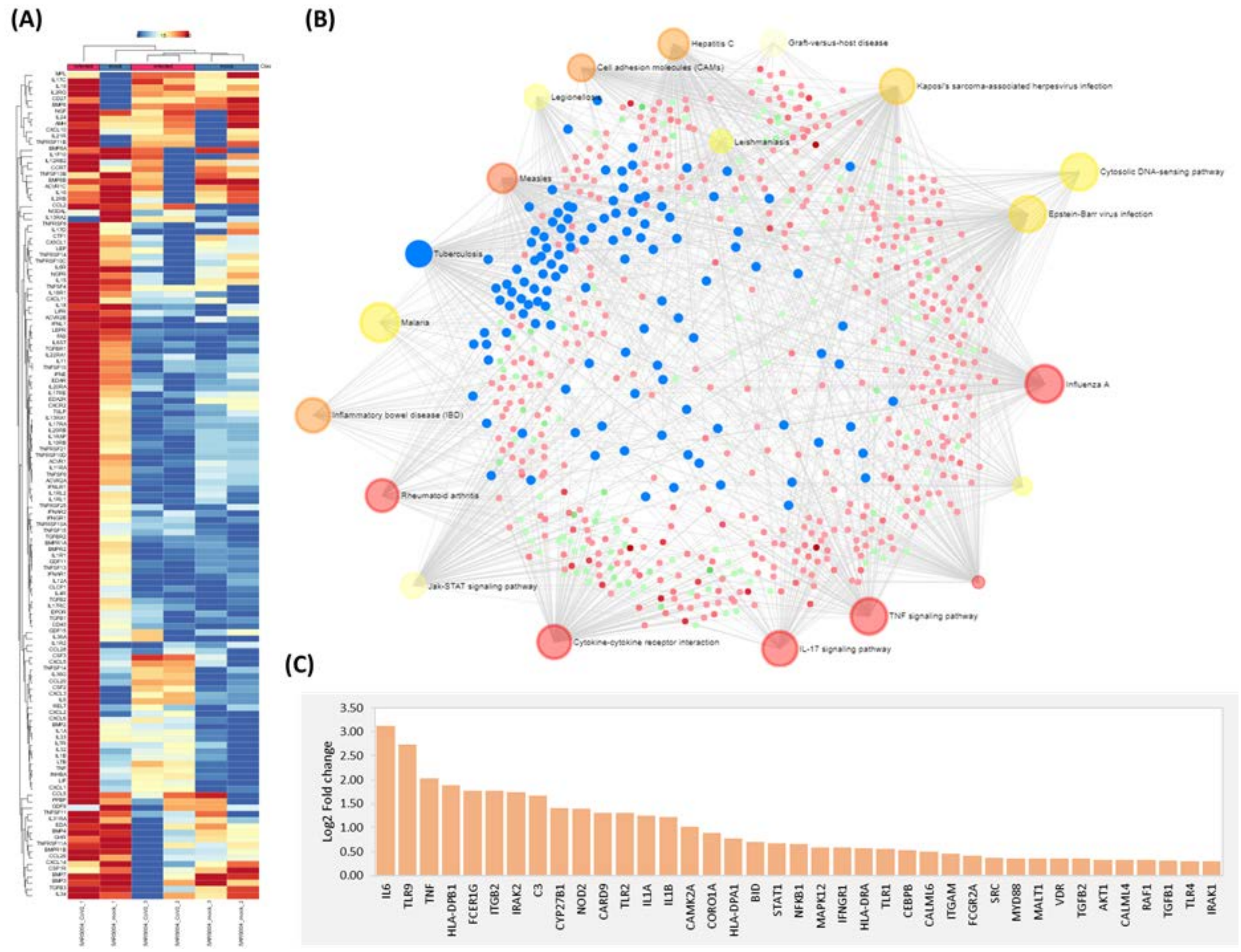


**Supplementary Figure S2. Gene set enrichment analysis in Tuberculosis pathway (KEGG ID: hsa05152).** (A) Expression intensity of genes in the Tuberculosis KEGG pathway in three SARS-CoV-2 infected cells and mock-treated cells. (B) Enrichment of differentially expressed genes (blue) in Tuberculosis pathway. (C) Fold change of up-regulated TB-related genes following SARS-CoV-2 infection compared to mock-treated cell lines.

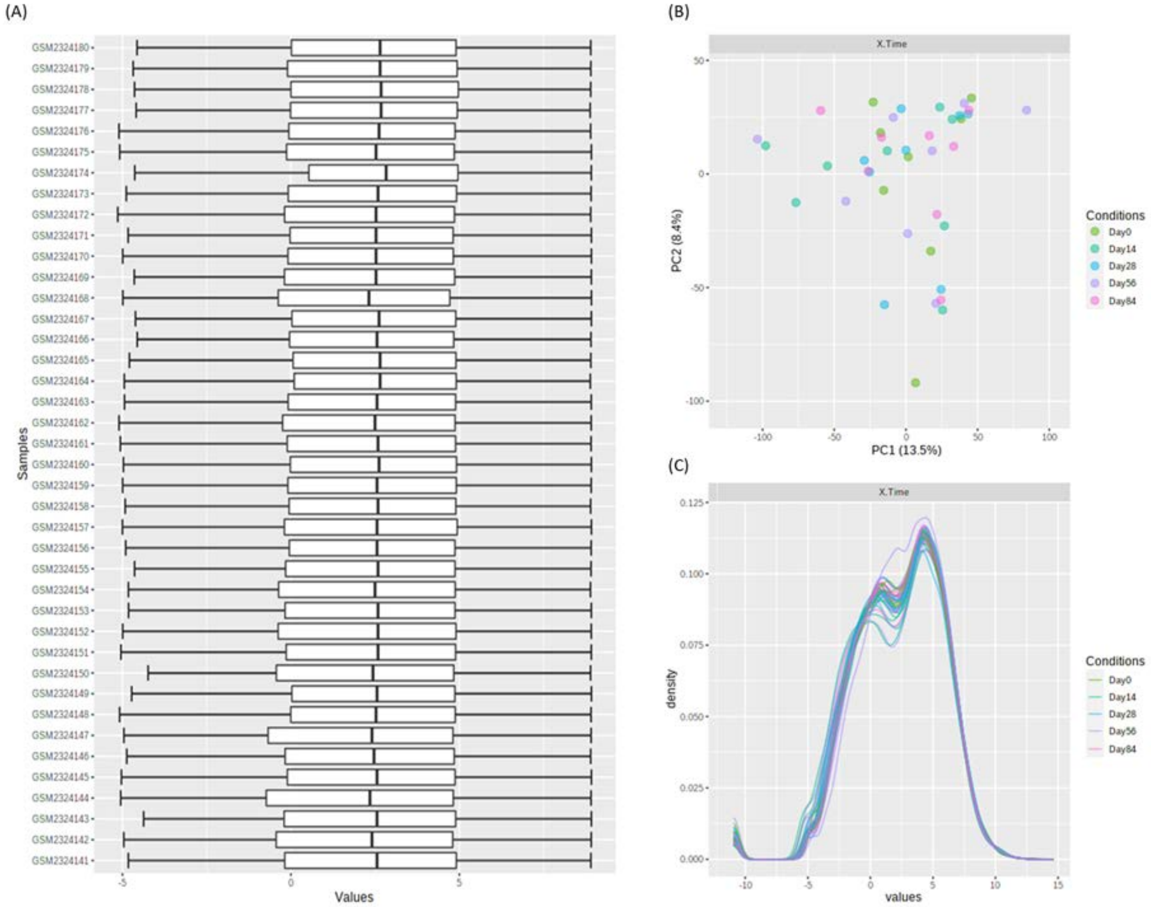


**Supplementary Fig. S3. Data exploration and quality check after normalization of RNA seq data (GSE87186).** (A) Box plot for the 40 samples included in the dataset. (B) Principal component analysis showing variations in the data. Each point represents a sample. (C) Density plot showing uniform distribution between samples.

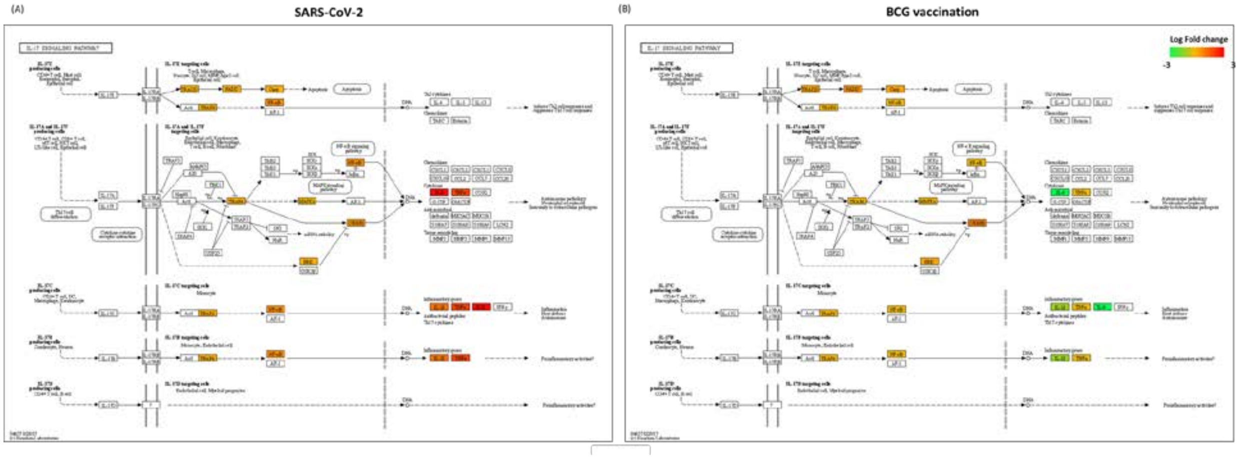


**Supplementary Figure S4. Expression level of DEGs in Interleukin 17 signaling pathway.** Colored by the log fold change of DEGs (A) DEGs in SARS-CoV-2 infection compared to mock-treated cells, (B) DEGs following BCG vaccination.

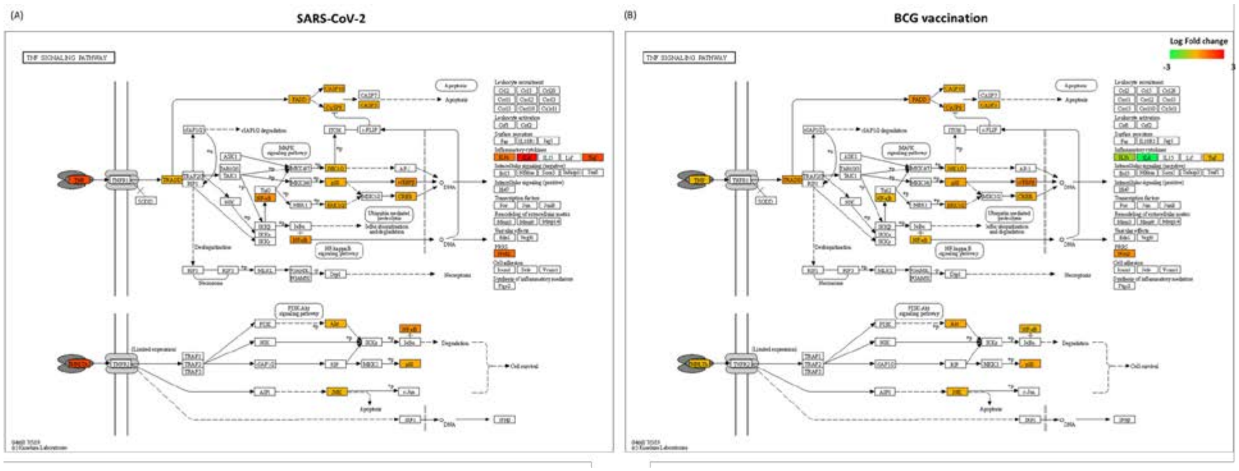


**Supplementary Figure S5. Expression level of DEGs in TNF signaling pathway.** Colored by the log fold change of DEGs (A) DEGs in SARS-CoV-2 infection compared to mock-treated cells, (B) DEGs following BCG vaccination.

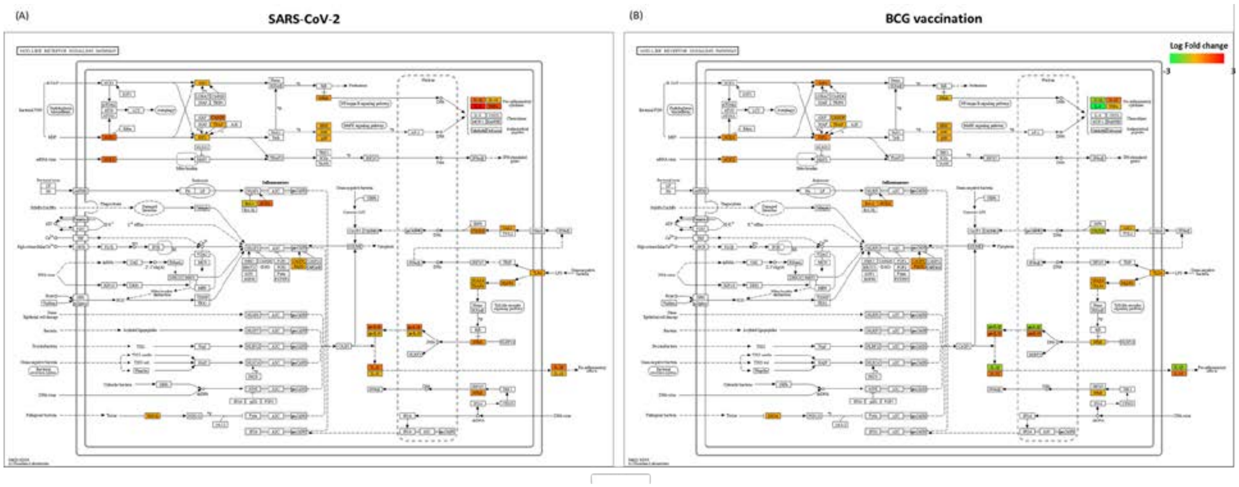


**Supplementary Figure S6. Expression level of DEGs in NOD-like receptor signaling pathway.** Colored by the log fold change of DEGs (A) DEGs in SARS-CoV-2 infection compared to mock-treated cells, (B) DEGs following BCG vaccination.

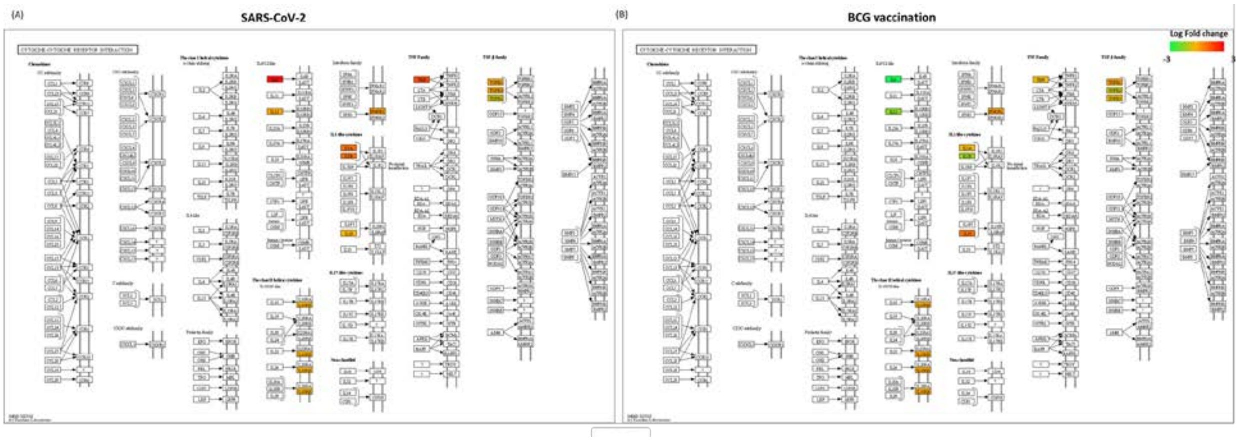


**Supplementary Figure S7. Expression level of DEGs in cytokine-cytokine receptor interaction pathway.** Colored by the log fold change of DEGs (A) DEGs in SARS-CoV-2 infection compared to mock-treated cells, (B) DEGs following BCG vaccination.

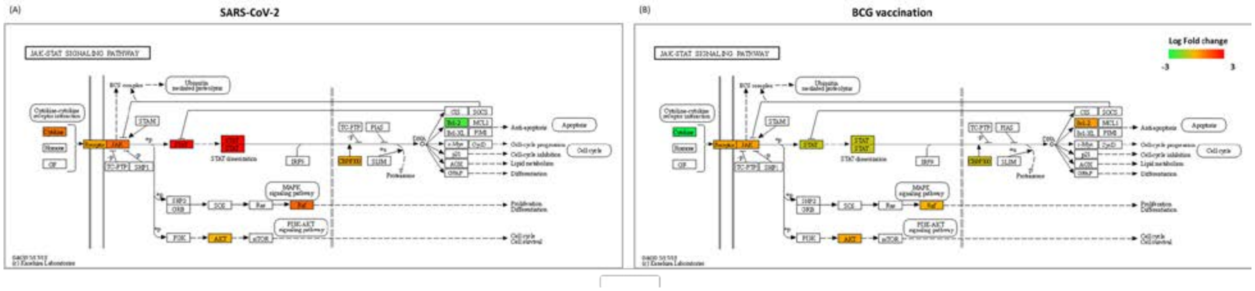

**Supplementary Figure S8. Expression level of DEGs in JAK-STAT signaling pathway.** Colored by the log fold change of DEGs (A) DEGs in SARS-CoV-2 infection compared to mock-treated cells, (B) DEGs following BCG vaccination.

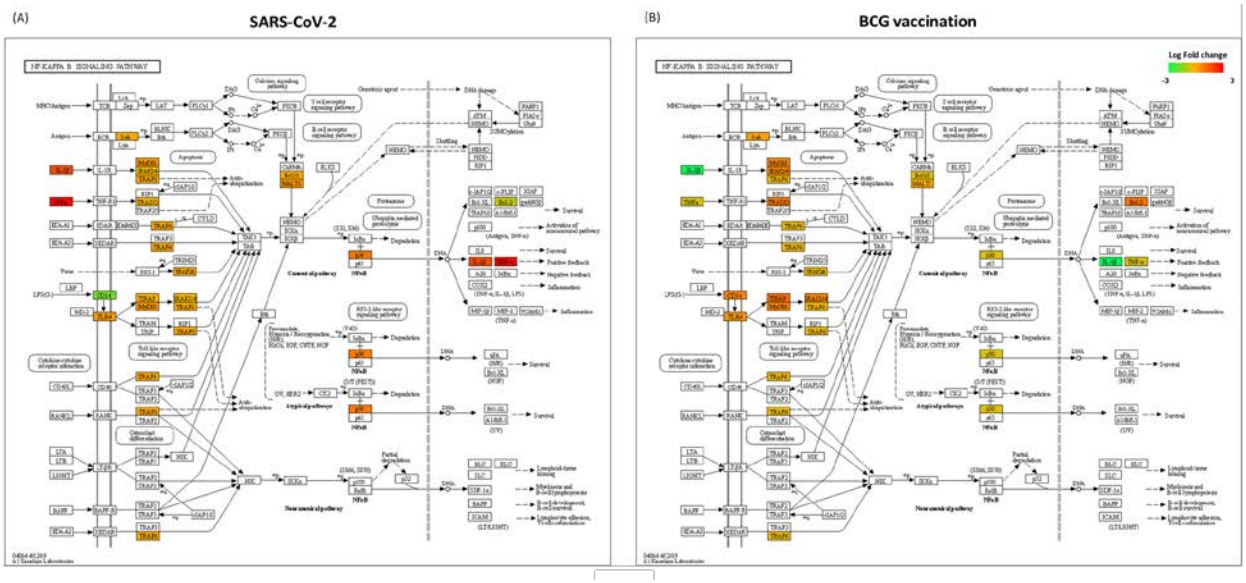


**Supplementary Figure S9. Expression level of DEGs in NF-Kappa B signaling pathway.** Colored by the log fold change of DEGs (A) DEGs in SARS-CoV-2 infection compared to mock-treated cells, (B) DEGs following BCG vaccination.

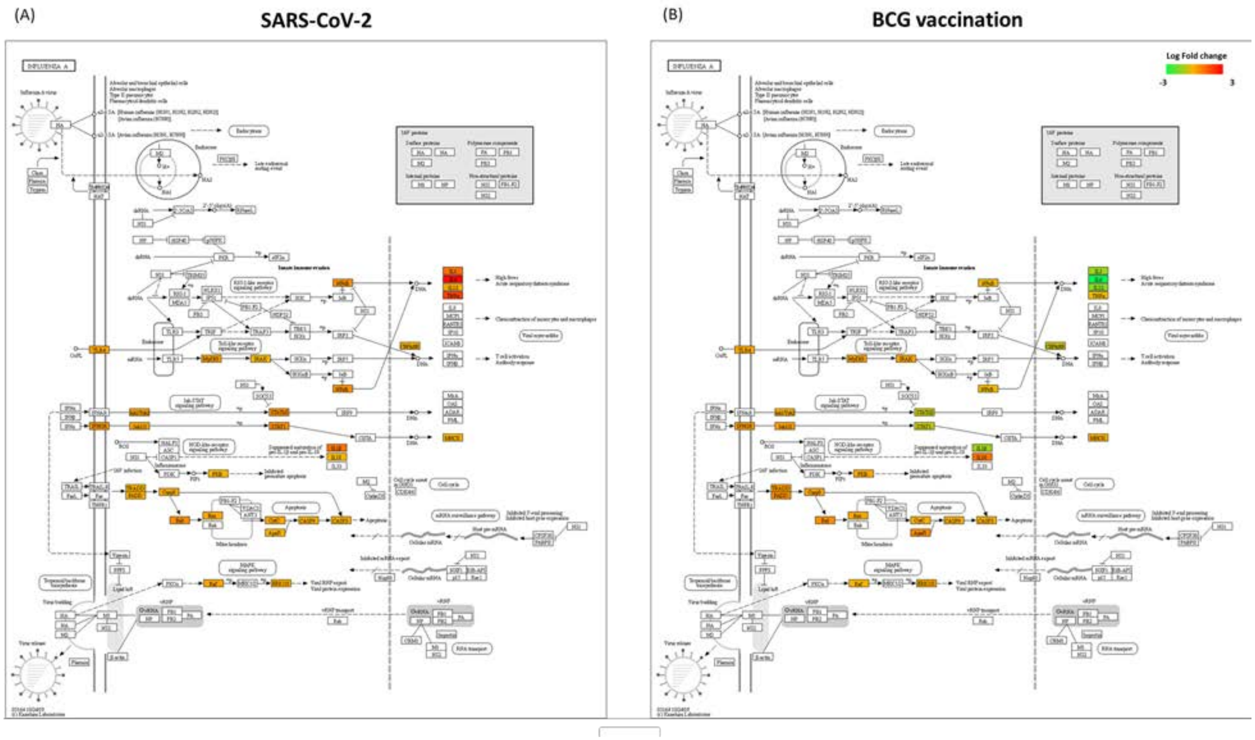

**Supplementary Figure S10. Expression level of DEGs in Influenza A pathway.** Colored by the log fold change of DEGs (A) DEGs in SARS-CoV-2 infection compared to mock-treated cells, (B) DEGs following BCG vaccination.
